# A low-cost smartphone fluorescence microscope for research, life science education, and STEM outreach

**DOI:** 10.1101/2022.07.06.498871

**Authors:** Madison A Schaefer, Heather N Nelson, John L Butrum, James R Gronseth, Jacob H Hines

## Abstract

Much of our understanding of cell and tissue development, structure, and function stems from fluorescence microscopy. The acquisition of colorful and glowing images engages and excites users ranging from seasoned microscopists to STEM students. Fluorescence microscopes range in cost from several thousand to several hundred thousand US dollars. Therefore, the use of fluorescence microscopy is typically limited to well-funded institutions and biotechnology companies, research core facilities, and medical laboratories, but is financially impractical at many universities and colleges, primary and secondary schools (K-12), and in science outreach settings. In this study, we developed and characterized components that when used in combination with a smartphone or tablet, perform fluorescence microscopy at a cost of less than $50 US dollars per unit. We re-purposed recreational LED flashlights and theater stage lighting filters to enable viewing of green and red fluorophores including EGFP, DsRed, mRFP, and mCherry on a simple-to-build frame made of wood and plexiglass. These devices, which we refer to as glowscopes, were capable of 10 μm resolution, imaging fluorescence in live specimens, and were compatible with all smartphone and tablet models we tested. In comparison to scientific-grade fluorescence microscopes, glowscopes may have limitations to sensitivity needed to detect dim fluorescence and the inability to resolve subcellular structures. We demonstrate capability of viewing fluorescence within zebrafish embryos, including heart rate, rhythmicity, and regional anatomy of the central nervous system. Due to the low cost of individual glowscope units, we anticipate this device can help to equip K-12, undergraduate, and science outreach classrooms with fleets of fluorescence microscopes that can engage students with hands-on learning activities.

## Introduction

Advances in smartphone and tablet camera technology have placed powerful cameras in the hands of scientists, educators, and students. The resolution and sensitivity of modern smartphone and tablet cameras surpasses the capabilities of many scientific cameras still being used for research applications. With annual improvements to smartphone camera sensitivity, pixel binning and resolution, and video frame rate capabilities, we are entering an era where these devices can perform high quality image and video acquisition for science education and research. Accordingly, products have been developed to mount smartphones onto microscope eyepieces for image acquisition. In addition, standalone ‘smartphone microscope’ designs have been developed for use in K-12 and undergraduate science education settings ^1 2 3^, research or clinical settings ^4-7^, and geoscience applications ^8^.

Fluorescence microscopy produces visually attractive images that can greatly enhance the contrast of specific features of the specimen being viewed. This microscopic imaging modality has historically involved costly and cumbersome mercury arc lamps or lasers. However, in recent years, comparatively inexpensive and compact laser-emitting diode (LED) technology is replacing mercury arc lamps and lasers in the microscopy industry. Advantages include lower cost, greater longevity, and maintenance free operation. Because LEDs have been engineered to emit light of virtually any wavelength on the visible light spectrum, and can be filtered if needed, the microscopy community is widely adopting use of LEDs as an excitation light source for fluorescence microscopy.

In this study, we aimed to develop a smartphone fluorescence imaging setup with a target build price of under $50 (US dollars) per unit. We demonstrate the ability of these devices, which we refer to as ‘glowscopes’, to detect green and red fluorophores and to monitor and detect changes to heart rate and rhythmicity in embryonic zebrafish. We discuss and demonstrate opportunities for undergraduate and K12 educators to use these resources to meet local and Next Generation Science Standards.

## Methods

### Animal care and use

All zebrafish work performed in this study was approved by the Institutional Animal Care and Use Committee at Winona State University. Zebrafish (Danio rerio) embryos were raised at 28.5**°**C in egg water (0.0623 g/L Coralife marine salt) and staged according to hours post-fertilization or morphological criteria. Transgenic lines used in this study included *Tg(olig2:DsRed)*^*vu19*^, *Tg(nkx2*.*2a:EGFP-CaaX)*^*vu16*^, *Tg(phox2bb:EGFP)* ^*w37*^, *Tg(mbpa:EGFP-CaaX; myl7:mCherry)* (hereafter referred to as *Tg(myl7:mCherry)* ^9^, *Tg(sox10:mRFP)*^*vu234*^, and *Tg(UAS:EGFP-CaaX, myl7:EGFP)*^*co18*^, *which is hereafter referred to as Tg(myl7:EGFP)*. To reversibly paralyze embryos and larvae for viewing, Tricaine Methanesulfonate (Western Chemical, Inc.) stock solution (0.116 M) was added directly to the petri dish containing egg water at a ratio of 0.5 - 1 ml (tricaine) into 20 ml (egg water). An astemizole stock solution (10 mM in DMSO) was used to treat zebrafish by direct addition to the egg water for a final concentration of 10 or 30 μM (as indicated). All other animals used in this study were collected locally prior to use. Viewing of live insects contained in petri dishes was aided by placing the dish over ice. A ReptiTherm U.T.H. was used to increase temperature for heart rate studies (Zoomed). The surface of this heating pad was measured at 35ºC at the time of use using an infrared thermometer. Treatment of zebrafish embryos with acidic water was performed using household vinegar (10% in egg water) for 30 minutes between pre-and post-treatment determination of heart rate. For fish prey-induced behavior, a single zebrafish larva (5 dpf) was placed into a 30 mm petri dish containing 5 ml egg water. For viewing of zebrafish plus food (Paramecium bursaria, living; Carolina Biologicals) was pre-mixed directly to the egg water at a concentration of 30 paramecium per 1 ml, which was determined prior to their addition by microscopic viewing of triplicate samples and diluted to achieve this target concentration.

### Glowscope components, assembly, and determination of image resolution

A complete parts list, design, and assembly information is available in Supplementary Methods. Briefly, cut plywood or composite material were assembled with a plexiglass (polycarbonate or acrylic) surface. Holes were drilled into the plexiglass to provide a clear viewing port for the smartphone camera. An additional plexiglass piece was cut to serve as a moveable stage platform immobilized by washers and clamps. A clip-on macro lens (Lieront, purchased from www.amazon.com, advertised as 25X macro) was used to add magnification to smartphone images for fluorescence. A USAF 1951 Test Chart (www.edmundoptics.com, Catalog #3857) was used to determine the maximum image resolution achieved. After the image was acquired with maximal digital zoom, we determined the smallest lines that could be clearly separated and the lp/mm they represent (x). Resolution (in microns) was calculated as follows: μm= 1000 / (x * 2).

### Glowscope fluorescence

A blue LED headlamp (Topme, purchased from www.amazon.com, advertised as a fishing headlamp) or a multi-color LED flashlight (Lumenshooter, purchased from www.amazon.com, advertised as a tactical flashlight) was used for GFP illumination. To reduce stray light in the green and yellow wavelengths, theater stage gel lighting film (Rosco #4990, CalColor Lavender) was cut and inserted between the LED and the focusing lens of the headlamp. Rosco #14 (Medium Straw) and #312 (Canary) theater stage lighting gel film were used individually or in combination as emission filters. The LED lamp was held at approximately a 45 degree angle above and within 3-6 inches of the specimens. Emission filters were placed between the acrylic platform and the clip-on lens. For imaging of red fluorescent proteins, Rosco #88 (Light Green) and #89 (Moss Green) were used for illumination (excitation filters) along with #19 (Fire) as an emission filter. All Rosco filters were purchased from purchased from www.stagelightingstore.com or www.bhphotovideo.com. A Vernier Emissions Spectrometer (VSP-EM, www.vernier.com) was used to assess the effects of excitation filters. Vernier Spectral Analysis software (version 4.11.0-1543) was used to acquire and convert data to .csv format, which was them imported into Microsoft Excel and subsequently to make plots using GraphPad Prism (version 7.0, GraphPad).

### Video acquisition and analysis

Apple and Samsung devices were used for image acquisition. Unless otherwise noted, all image acquisition was performed using an iPhone XR (Apple). For convenience, time-lapse videos of the embryonic zebrafish heart were acquired using the ProCam 8 app, which allowed the 1X lens to be locked for use, and digital zoom extended to 4.0X. Video acquisition at 1080p resolution and 60 frames per second (fps) was used because use 4k resolution or higher frame rates decreased fluorescence sensitivity and signal:noise ratio. All images and videos were transferred to a laptop using Airdrop (Apple) or a USB cable to avoid video data compression. Videos were imported into Fiji ^10^ by first converting videos to a series of TIFF images (one per frame) using Adobe Photoshop. After opening smartphone videos in Photoshop, images were cropped and then saved as an image sequence using the File → Export → Render Video → Photoshop Image Sequence (TIFF format) command. The image sequence was then opened in Fiji through the File → Import → Image Sequence command. For non-fluorescent time-lapse viewing of paramecia, zebrafish swimming, and tadpole and caterpillar escape responses, we used 1080p resolution with 120 fps acquisition.

For zebrafish heart videos, detection and measurement of heart chamber movements in some videos was aided by edge detection. In these instances, Fiji was used to detect edges using the Process → Find Edges command. To detect fluorescence changes associated with atrium and ventricle movements, the raw or edge-detected image sequence was used and further analyzed within Fiji. First, the Analyze → Set Measurements command was used to instruct Fiji to measure minimum and maximum gray values, mean gray values, and limit to threshold. The image stack was converted to greyscale using the Image → Type → 8-bit command. In instances where image drift or non-biological movements occurred, the Plugins → Image Stabilizer command was used in attempt to eliminate this issue. The image was scanned for potential regions of interest (ROI) where the heart chamber walls consistently moved back and forth across the x-y axis. Once a ROI was defined, it was added using the Analyze → Tools → ROI manager window or the “t” shortcut. Concurrently to identifying a ROI, the Image → Adjust → Threshold window was used to establish a threshold whereby the chamber walls repeatedly occupied and vacated ROI box. Once an ideal combination of image threshold and ROI box position was established, the image was converted to binary data using the Process → Binary command selecting default, dark, and black background settings. To measure fluorescence intensity within the ROI over time, Multi Measure tool within the ROI manager (under ‘More’ tab) was used to create a set of measurements over time. Raw data were cut and pasted directly into Microsoft Excel spreadsheet. Within the spreadsheet, formulas were used to identify the onset of each heart beat using the maximum intensity column and plots were made using GraphPad Prism 7.0. The onset of each beat was defined by the transition from 0 to 255.

## Results

### Development of a low-cost smartphone fluorescence microscope

As a frame to support a smartphone or tablet for fluorescence viewing, we adapted a do-it-yourself microscope design used widely for non-fluorescent imaging ^3^. This frame consisted of a plexiglass platform to hold a smartphone (or tablet) atop the specimen (Fig. 1A). A second plexiglass platform positioned below the smartphone lens served as a stage to hold the specimen. The adjustable height of the plexiglass stage ensured the ability to focus on the specimen. Wood or composite boards, nuts, bolts, and washers held the plexiglass platform and stage in place. For non-fluorescent imaging, a battery-powered LED work lamp positioned below the specimen provided additional light when ambient room lighting was insufficient (Fig. 1A).

**Figure 1.**
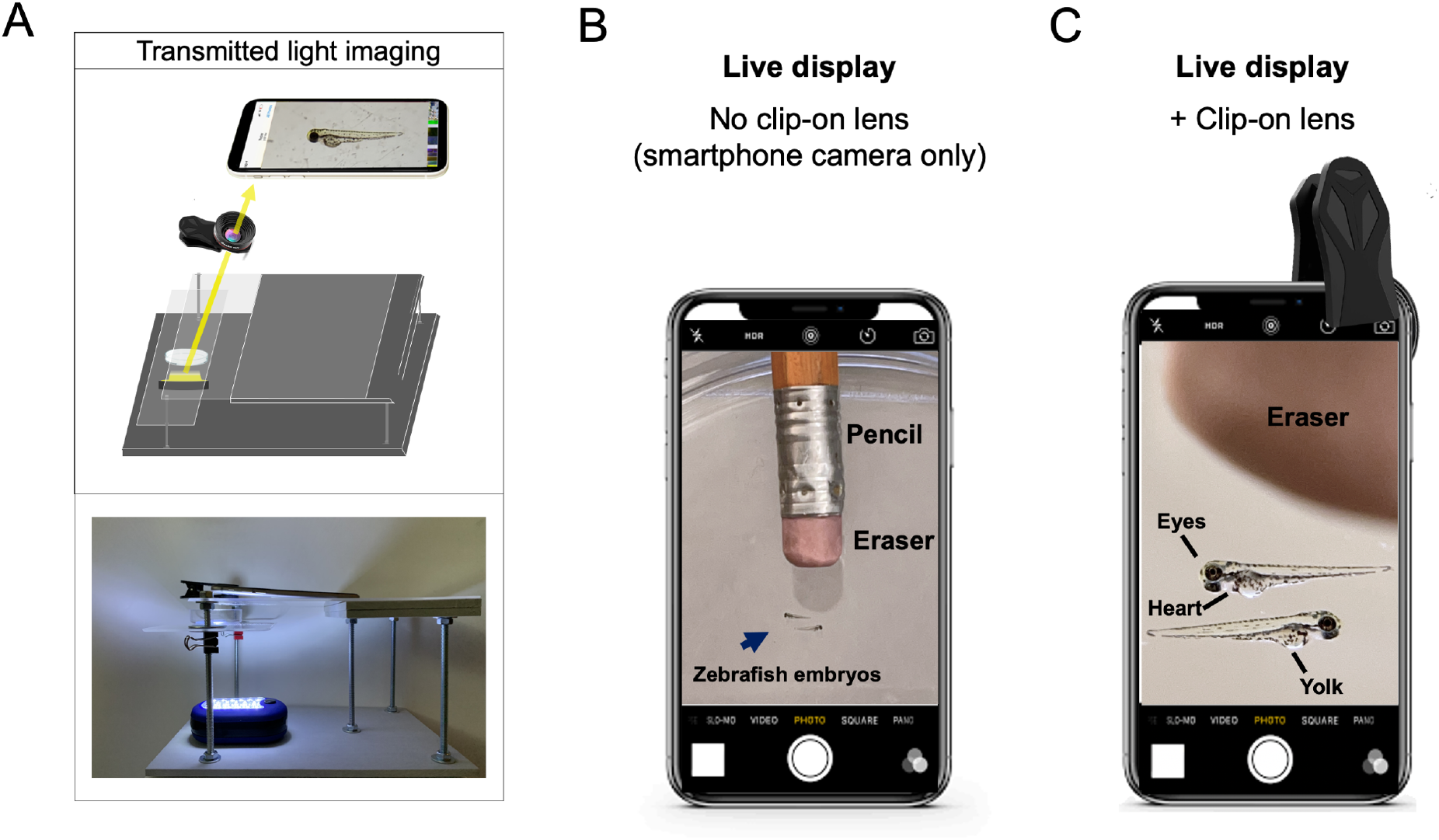
Live view of non-fluorescent specimens using the glow scope frame. (**A**) Illustration shows the light path for transmitted light viewing. Components shown include the smartphone, clip-on lens (black), glowscope frame (gray), stage and viewing platform (transparent), petri dish containing a specimen, and a LED work lamp positioned under the stage and viewing platform. (**B**) Image view using a smartphone camera without the clip-on lens shows zebrafish embryos (blue arrow). A standard pencil (eraser side) is shown as a size reference. (**C**) Image view using the additional clip-on lens shows the same zebrafish embryos and pencil eraser seen in (B). Images shown in B-C are native magnification (not stretched) as seen on live-view, acquired using an Apple iPhone 12 Pro with 1x (middle magnification) lens, 6x digital zoom.

To increase magnification and resolution beyond the smartphone or tablet alone, an additional ‘clip-on’ macro lens was clamped over the phone or tablet camera (Fig. 1A, C). The additional clip-on lens used in this study provided approximately 5-fold magnification (Fig. 1B-C). This magnitude depended on the distance the specimen was placed from the clip-on lens. When viewing embryonic zebrafish (2-4 days post-fertilization), which are roughly equivalent to the length of a pencil eraser (2-3 mm in length), the increased magnification and image resolution made features of the zebrafish body plan easily observed. For example, anatomical structures including the eye, heart, yolk, and tail were easily observed on the live display using the clip-on lens (Fig. 1C). Using USAF test targets, we measured 9.8 μm resolution using an iPhone 11 camera with the added clip-on lens. This corresponded to significant improvements to image resolution and anatomical detail in live view or after image acquisition. After images were captured and digitally stretched to further magnify on a smartphone or laptop screen, the clip-on lens easily resolved anatomical structures and even individual pigment cells (melanocytes). In comparison, without the clip-on lens, these same features were difficult or impossible to view (Fig. 2)

**Figure 2.**
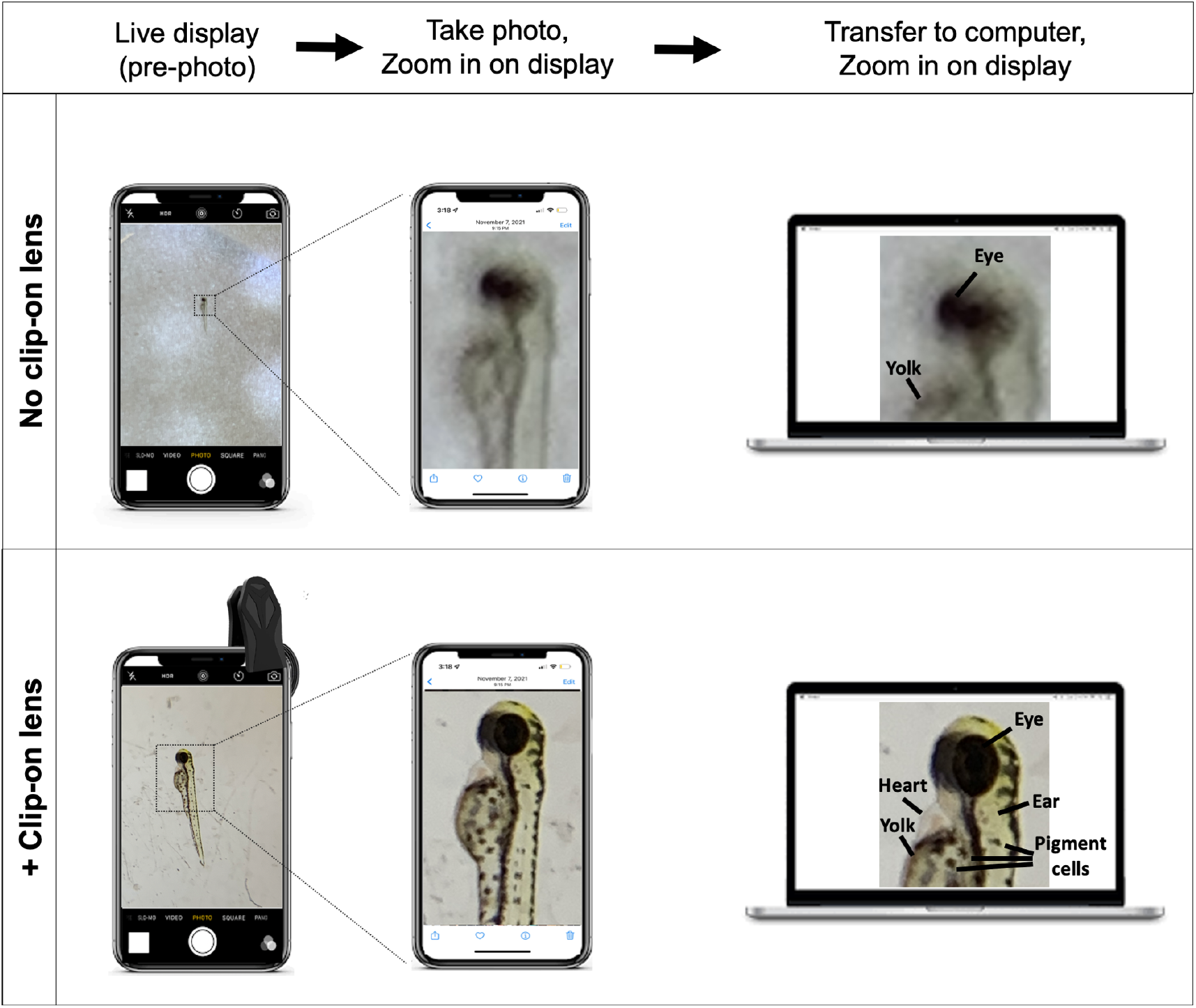
Photo capture and image stretching provides further magnification and anatomical detail. All images show 3 dpf zebrafish embryos (lateral view). Upper and lower images show the effect of taking and zooming in on photos with and without the clip-on lens, respectively. Note the benefits of digital magnification (image stretching) after image capture were most beneficial when the clip-on lens is used. Images were acquired using an Apple iPhone 12 Pro with 1x (middle magnification) lens, 6x digital zoom.

To equip the smartphone scope with fluorescence imaging capability, we next added fluorescence light sources and filters. To view specimens with green fluorescence, we turned off the LED work light and instead used a blue LED headlamp or flashlight (Fig. 3A). We used theater stage lighting filters to block the blue light from reaching the camera lens. To do so, we placed the filter immediately underneath the clip-on lens and smartphone (Fig. 3A). These inexpensive ‘emission filters’ reduced fluorescence background from the blue flashlight while permitting the green fluorescence from the specimen to pass through the filter and reach the camera. We initially obtained a sample-pack of theater stage lighting filters with a wide range of spectral properties. After testing numerous filter color options, we found that the combination of Rosco #14 and #312 stage-lighting filters effectively blocked blue light while enabling robust green fluorescence signal to transmit through to the camera (Fig. 3A). To test the functionality of this setup, we used transgenic zebrafish reporter lines that expressed enhanced green fluorescent protein (EGFP) in subsets of tissues or neural cells. For example, the fluorescence setup easily detected brain and spinal cord fluorescence in *Tg(nkx2*.*2a:memGFP)* embryos, cardiac tissue in *Tg(myl7:EGFP)* embryos, and had adequate sensitivity to detect EGFP expressed in a sub-region of the embryonic hindbrain in *Tg(phox2b:EGFP)* embryos (Fig. 3B-D). To further reduce background fluorescence and enhance the signal:noise ratio, we added Rosco filter 4990 positioned between the blue LED flashlight and the specimen (Fig. 3A). Anecdotally, this ‘excitation’ filter darkened the otherwise blue-green background with negligible loss of green fluorescence emitted and transmitted to the smartphone camera and viewed image. Using a Vernier spectrometer, we found that the added excitation filter slightly reduced the intensity of LED fluorescence between 460 – 500 nm (Fig. 3F-G). This may explain the reduced blue background fluorescence on the viewed images when R4990 excitation filter was used. Though this additional excitation filter is not necessary for green fluorescence viewing, it can effectively reduce background and improve the signal:noise ratio when viewing specimens with dim fluorescence.

**Figure 3.**
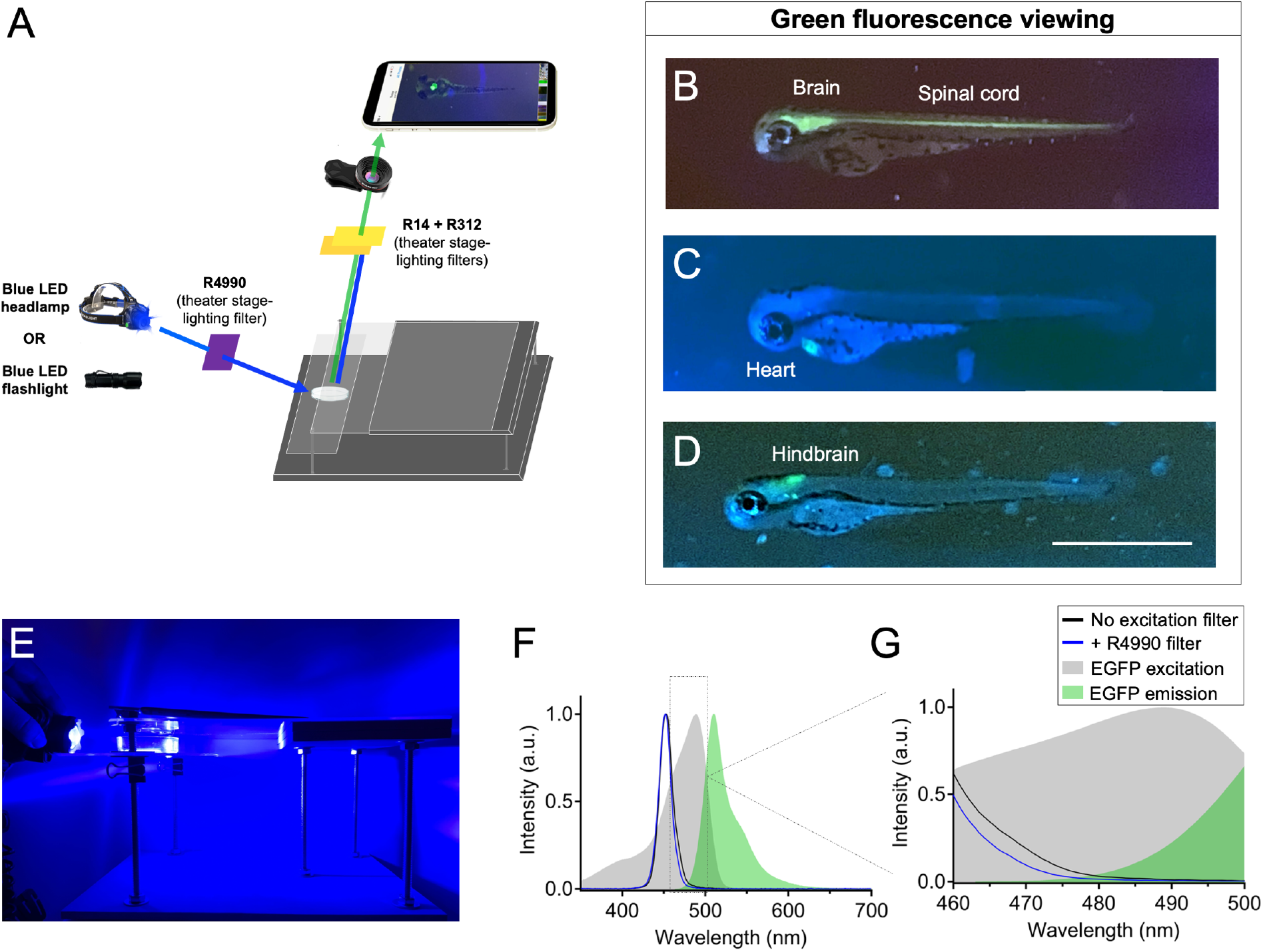
Use of recreational LED flashlights and theater stage lighting filters for smartphone green fluorescence microscopy. (**A**) Schematic of components used for green fluorescence viewing on the glowscope. (**B-D**) Representative fluorescence images of transgenic zebrafish embryos (3-4 dpf, lateral views) expressing green fluorescence in cell-type specific patterns. Reporter lines viewed include *Tg(nkx2*.*2a:memGFP)* (B), *Tg(myl7:EGFP)* (C), *and Tg(phox2b:EGFP)* (D). (**E**) Image shows the glowscope in use for green fluorescence viewing. (**F-G**) Plots show the blue LED flashlight emission wavelength (black and blue lines, measured using a Vernier spectrometer) in comparison to the excitation profile of EGFP (gray, obtained from www.fpbase.org). The dashed box region in F is further magnified in G to show the effect of the R4990 filter with greater detail. Images in B-D were acquired using an Apple iPhone XR.

We also tested LEDs and stage lighting filters for viewing red fluorescence (Fig. 4). In place of a blue flashlight we instead used a green LED flashlight. As an emission filter positioned underneath the lens, we used Rosco stage lighting filter #19 to block green light while permitting red fluorescence to pass through to the camera lens (Fig. 4A). In our hands, this emission filter worked best when doubled up, and provided clear fluorescence viewing of transgenic zebrafish lines expressing red fluorescent proteins. As examples, this setup effectively illuminated brain and spinal cord features of *Tg(olig2:DsRed)* embryos, heart fluorescence in *Tg(myl7:mCherry)* embryos, and head and jaw skeleton (cranial neural crest) fluorescence in *Tg(sox10:mRFP)* embryos (Fig. 4B-D). The LED flashlight we used advertised 530 nm LED excitation, which among the fluorophores we used, aligned best with the spectral properties DsRed. In our hands, the green flashlight and Rosco stage lighting filters in fact paired best with DsRed, but somewhat less well with the further red-shifted fluorescent proteins mRFP and mCherry (Fig. 4B-D). Background fluorescence and signal:noise ratio were marginally improved by covering the flashlight with Rosco filters #88 and #89 (Fig. 4A). This excitation filtering reduced fluorescence intensity at wavelengths between 540 – 580 nm (Fig 5F-G), and may explain the decreased background fluorescence observed in acquired images when the excitation filters were used. Although the excitation filters #88 and #89 improved image quality and detection of dim fluorescence, they were not necessary for red fluorescence imaging in our hands.

**Figure 4.**
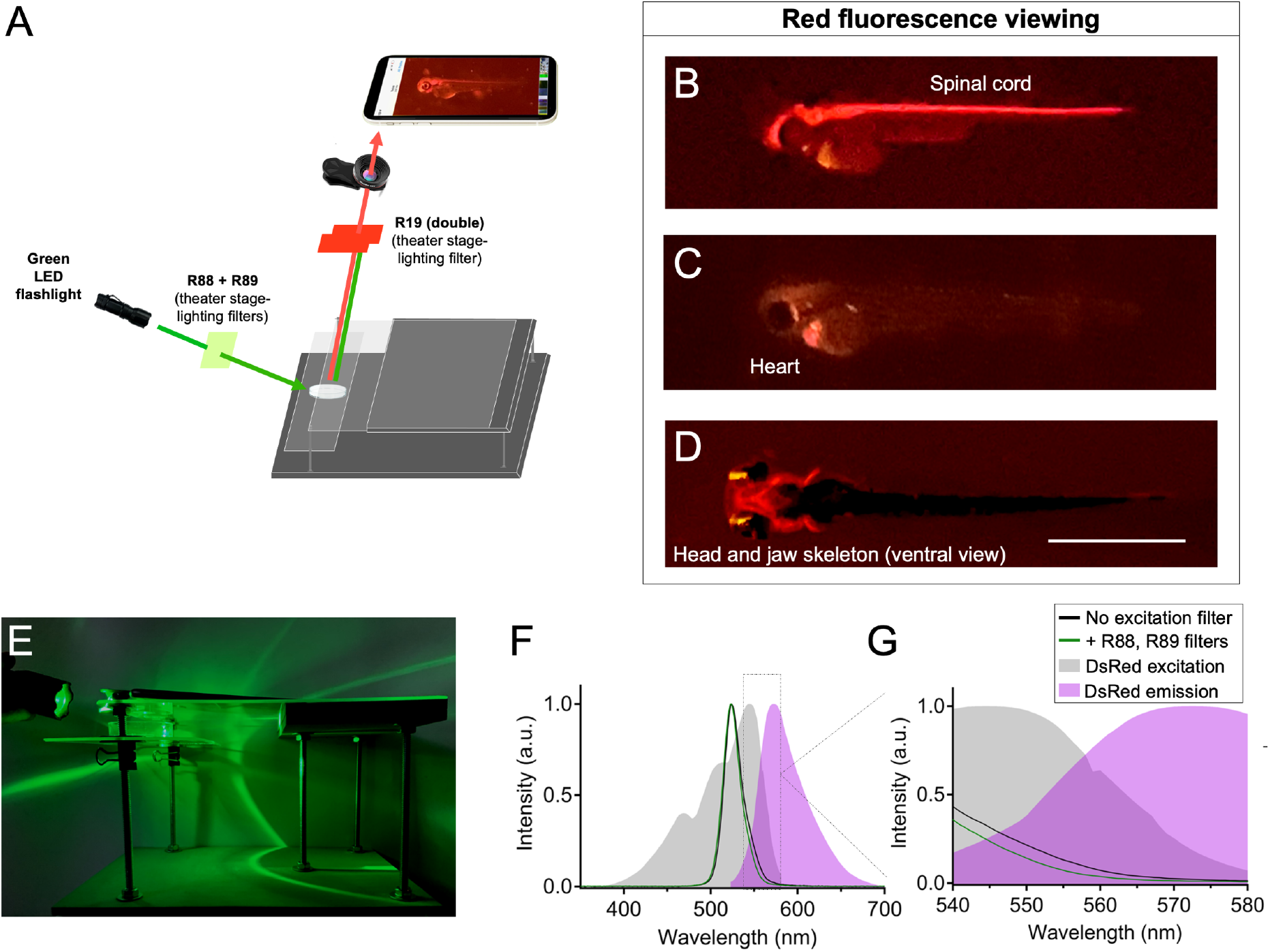
Glowscope detection of red fluorescent proteins. (**A**) Schematic of components used for red fluorescence viewing on the glowscope. In comparison to green fluorescence viewing, flashlight and filters are changed, but the remainder of the glowscope is unchanged. (**B-D**) Representative fluorescence images of transgenic zebrafish embryos (3-4 dpf, lateral views) expressing red fluorescence in cell-type specific patterns. Reporter lines viewed include *Tg(olig2:DsRed)* (B), *Tg(myl7:mCherry)* (C), *and Tg(sox10:mRFP)* (D). Scale bar is 1 mm. (**E**) Image shows the glowscope in use for red fluorescence viewing. (**F-G**) Plots show the green LED flashlight emission wavelength (black and green lines, measured using a Vernier spectrometer) in comparison to the excitation profile of DsRed (gray, obtained from www.fpbase.org). The dashed box region in F is further magnified in G to show the effect of the R88 + R89 filters with greater detail. Images in B-D were acquired using an Apple iPhone XR with the exception of D, which was acquired using an iPhone 12 Pro.

**Figure 5.**
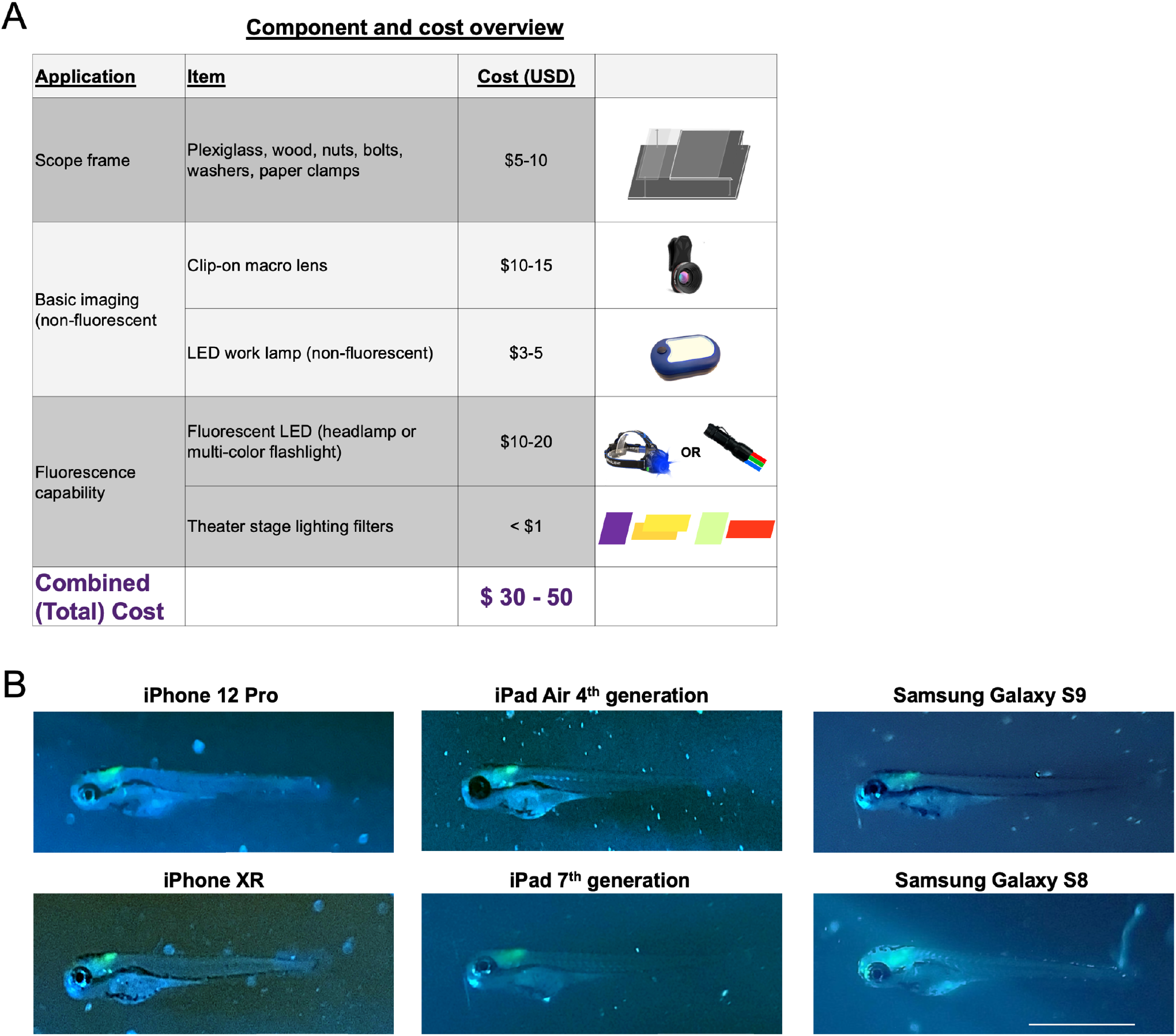
Summary of glowscope components and smartphone compatibility. (A) Table shows the basic components and costs (at the time of this study) in US Dollars. (B) Images of *Tg(phox2b:EGFP)* zebrafish embryos demonstrate compatibility with all devices tested, but subtle differences between various tablets and smartphones used for testing. Scale bar is 1 mm.

### Summary of glowscope components, cost, and smartphone compatibility

The parts needed to convert a smartphone or tablet into a fluorescence microscope for are summarized in Figure 5. At the time of our testing and manuscript preparation, the necessary parts were acquired for a total of $30-50 per individual glowscope (Fig. 5A). Wood and plexiglass were cut and drilled with basic tools. Theater stage lighting filters were purchased in large sheets and cut to size, resulting in a cost of less than $0.10 USD per unit filter for this application. Blue and green LED flashlights were readily available from online retailers as sold for tactical, hunting, and fishing applications. We tested numerous options and found that those with blue rather than UV light worked best for viewing green fluorescent protein. Our preferred light source was a multi-color tactical flashlight with both blue (454 nm) and green (530 nm) colors, which retailed for $20 USD at the time these studies were performed. By contacting sellers directly and purchasing in bulk, we arranged for direct purchasing at wholesale cost (∼50% discount).

To assess glowscope compatibility with different camera devices, we next tested the fluorescence viewing capabilities using various smartphone and tablet models to detect green fluorescence in *Tg(phox2b:EGFP)* embryos. Of the green reporter lines initially tested (Fig. 3B-D), *Tg(phox2b:EGFP)* was the most challenging to view because the green fluorescence expression is only present in part of the embryonic hindbrain. All devices tested, including Samsung and Apple phones and tablets, detected the subtle green fluorescence in the hindbrain (Fig. 5B). Newer devices generally acquired fluorescence images with brighter green intensities in comparison to the background fluorescence, but these differences were subtle. We conclude that use of the newest smartphone or tablet models is not necessary for glowscope fluorescence viewing.

### Glowscope determination of embryonic zebrafish heart rate and rhythmicity

We next tested the ability of the glowscope to detect changes to heart rate and rhythmicity using green fluorescence imaging in embryonic zebrafish. Because the *Tg(myl7:EGFP)* reporter expresses green fluorescence in cardiomyocytes, the glowscope and live smartphone display made the beating heart chamber movements clearly observable over time (Fig. 6A). To determine if the imaging setup was capable of detecting drug-induced changes to heart rate, we performed pre-and post-viewing before and after 30 min treatment with astemizole. Formerly branded as Hismanal, astemizole was commonly used as an antihistamine allergy medication, but was pulled from the market in 1999 because it caused rare but fatal heart complications such as long QT syndrome, cardiac arrythmias, and life-threatening tachycardia ^11^. These human side effects were caused because astemizole blocks ERG-type K^+^ channels involved in repolarizing the cardiac action potential ^12^ in addition to its antihistamine mode of action. Previous studies have demonstrated that astemizole treatment causes dose-and time-dependent bradycardia followed by eventual atrioventricular 2:1 block (arrythmia) and cardiac arrest in zebrafish embryos ^13^. Using smartphone fluorescence viewing of *Tg(myl7:EGFP)* zebrafish in real-time, we detected a significant reduction to heart rate following 30 min astemizole treatment (Fig 6B-C). This drug treatment-induced bradycardia was similarly observed while recording the video (live view), by observing the recorded video in playback on the smartphone display screen, or by transferring the video recording to view on a laptop display (Fig. 6D).

**Figure 6.**
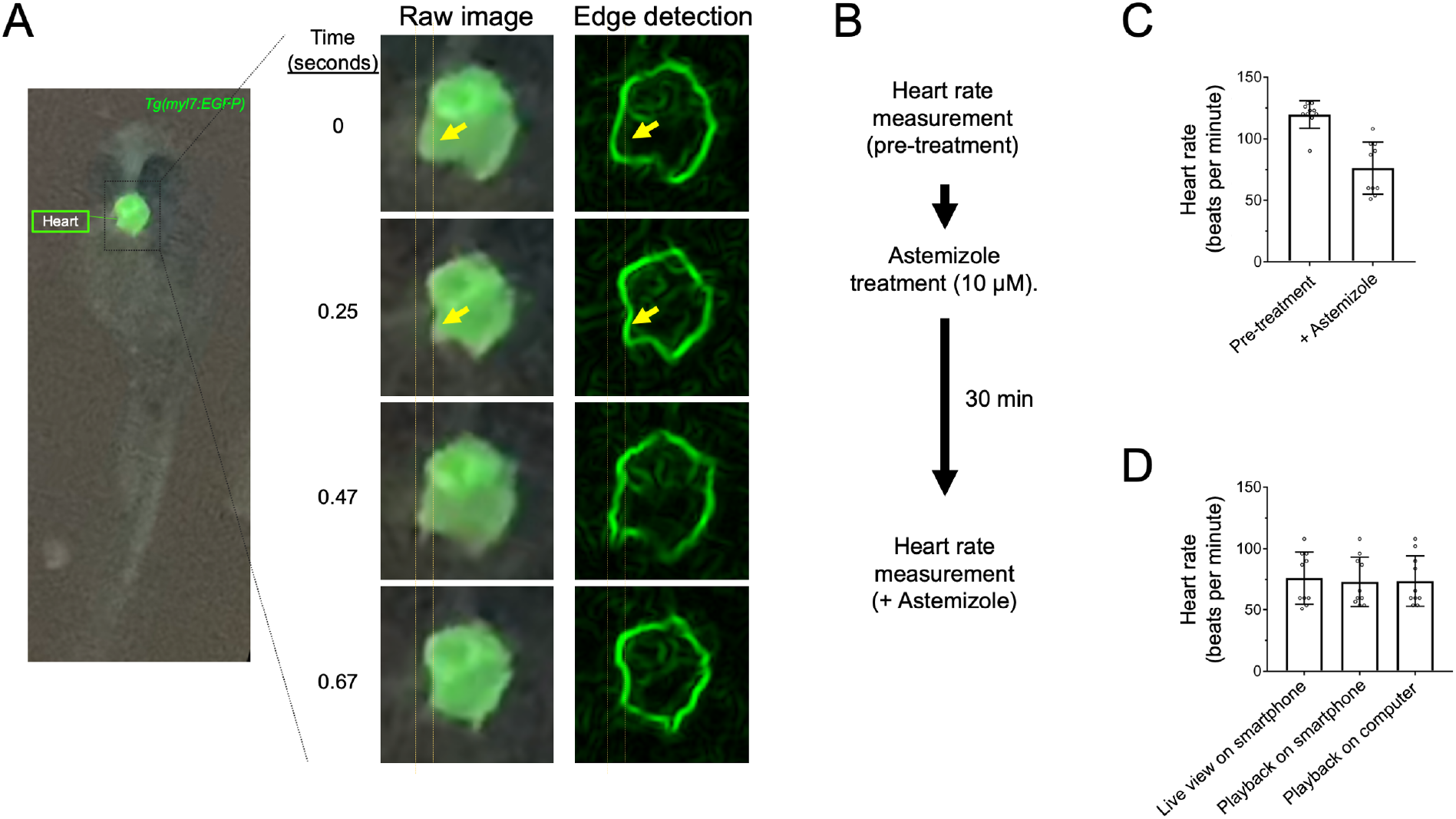
Glowscope detection of drug-induced changes to heart rate in transgenic zebrafish embryos. (**A**) Left fluorescence image shows GFP expression in the heart of a *Tg(myl7:EGFP)* embryo (ventral view). The boxed region is further magnified in the middle column, which from top to bottom show chamber movements across a video acquired on an iPhone 11 Pro smartphone. To aid viewing of chamber movements, these videos were transferred to a laptop, opened in ImageJ (free software), and processed using edge detection, which outlines the walls of the chambers. (**B**) Flow chart of the pre-and post-imaging of zebrafish heart rate in response to astemizole treatment (10 μM). (**C**) Plots show the average heart rate of zebrafish embryos prior to and after treatment with astemizole. (**D**) Summary of measurements conducted while recording the video (left bar) in real time, viewed on the smartphone after recording the video (middle bar), or viewed on a laptop (right bar). For C-D, Scatter plot points represent individual zebrafish (heart) rate measurements.

To further test the capabilities and limitations of smartphone fluorescence, we next assessed the ability to separately resolve atrium and ventricle chamber contractility rates, which we predicted could be beneficial to detect cardiac arrythmias. On most video recordings we captured, individual chamber movements were clearly visible on some but not all embryos, and pushed the limitations of the glowscope. To enhance the clarity and detection of chamber movements, we transferred the smartphone videos to a computer running the freely available software ImageJ (also known as Fiji). Within Fiji, the ‘find edges’ command provided a simple means to outline (edges of) the heart chambers and their movements over time (Fig. 7A-C). By monitoring fluorescence intensity changes specifically within a small region of interest (ROI) box, we detected oscillating heart chamber wall movements in and out of the ROI over time (Fig. 7C-D). This method provided clear separation of atrial and ventricular contractility rates using the smartphone device (Fig. 7D). Plots generated from videos recorded before and after astemizole treatment displayed notable differences. Prior to treatment, both chambers beat rapidly and in an alternating pattern. In comparison, 30-minute treatment with a high dose of astemizole (30 μM) caused severe bradycardia as well arrythmia, as evidenced by the ventricle stalling during every other atrium beat (Fig. 7D).

**Figure 7.**
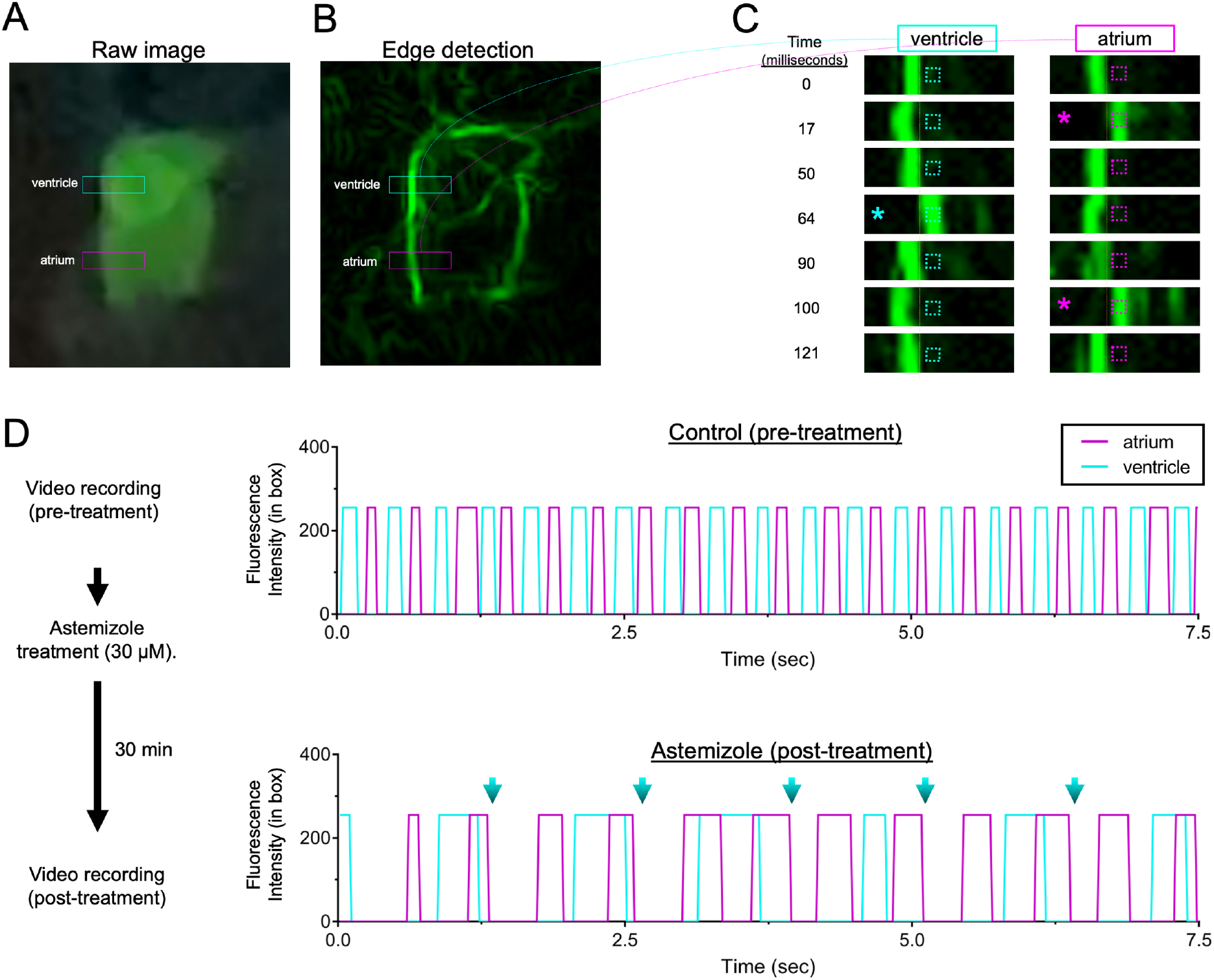
Glowscope detection of drug-induced cardiac arrythmia. (**A-B**) Still images from a glowscope video (acquired on an iPhone 11 Pro) show atrium and ventricle chambers in 3 dpf *Tg(myl7:EGFP)* zebrafish embryos. Images show either the raw (unedited) view of heart fluorescence directly from smartphone video recording (A) or a processed view of the same heart after edge detection was performed on a computer using Fiji (B). Blue and magenta rectangles show the regions of ventricle and atrium chambers used for high-magnification viewing and analysis in C-D. (**C**) Images (top to bottom) show a timecourse of ventricle (left) or atrium (right) chamber movements detected by the glowscope (corresponding to boxed regions in B). The small, dash-boxed regions of interest were used to monitor fluorescence intensity changes over time (**D**) Graphs of fluorescence intensity vs. time plotted for atrium and ventricle chamber reveals oscillating chamber movements. Measurements were separately acquired and analyzed for control (upper) and astemizole-treated (lower) embryos. The maximum fluorescence intensity (in arbitrary units) within the dash-boxed region of interest shown in panel C was obtained and plotted for each time frame of the video recording, resulting in a fluorescence peak each time the chamber fluorescence entered and occupied the boxed region of interest. Note the differences in both heart rate between control and astemizole-treated hearts as well as the arrhythmic 2:1 atrial:ventricular beat pattern induced by astemizole treatment (lower plot, 30 μM). Blue arrows in D show missed ventricular beats.

### Optimized lighting for non-fluorescent imaging

In addition to its use for fluorescence microscopy, the glowscope frame and clip-on lens can also be useful for viewing non-fluorescent specimens, and has been widely used in some science outreach settings ^3^. In our hands, a LED work lamp positioned underneath the specimen worked well for transparent specimens such as zebrafish embryos and larvae (Figs. 1, 2, 8A). Acknowledging that transmitted light may be ineffective for viewing non-transparent specimens, we next compared alternative lighting options for both transparent and non-transparent specimens. When viewing specimens such as tadpoles or insects, we found that positioning the light beside or above (epi-illumination) the specimen was advantageous (Fig. 8B-C). Either option could be achieved by holding the LED work lamp at the desired position by hand. As a more convenient option for epi-illumination, we re-purposed USB-powered COB LED strip lights. These inexpensive lights are normally sold for household cabinet and closet lighting and have adhesive (tape) backing, which enabled us to attach them under the plexiglass immediately atop the specimen (Fig. 8C). These alternative lighting options may be useful when specimen-type versatility is desired but were not necessary for our viewing of embryonic and larval zebrafish.

**Figure 8.**
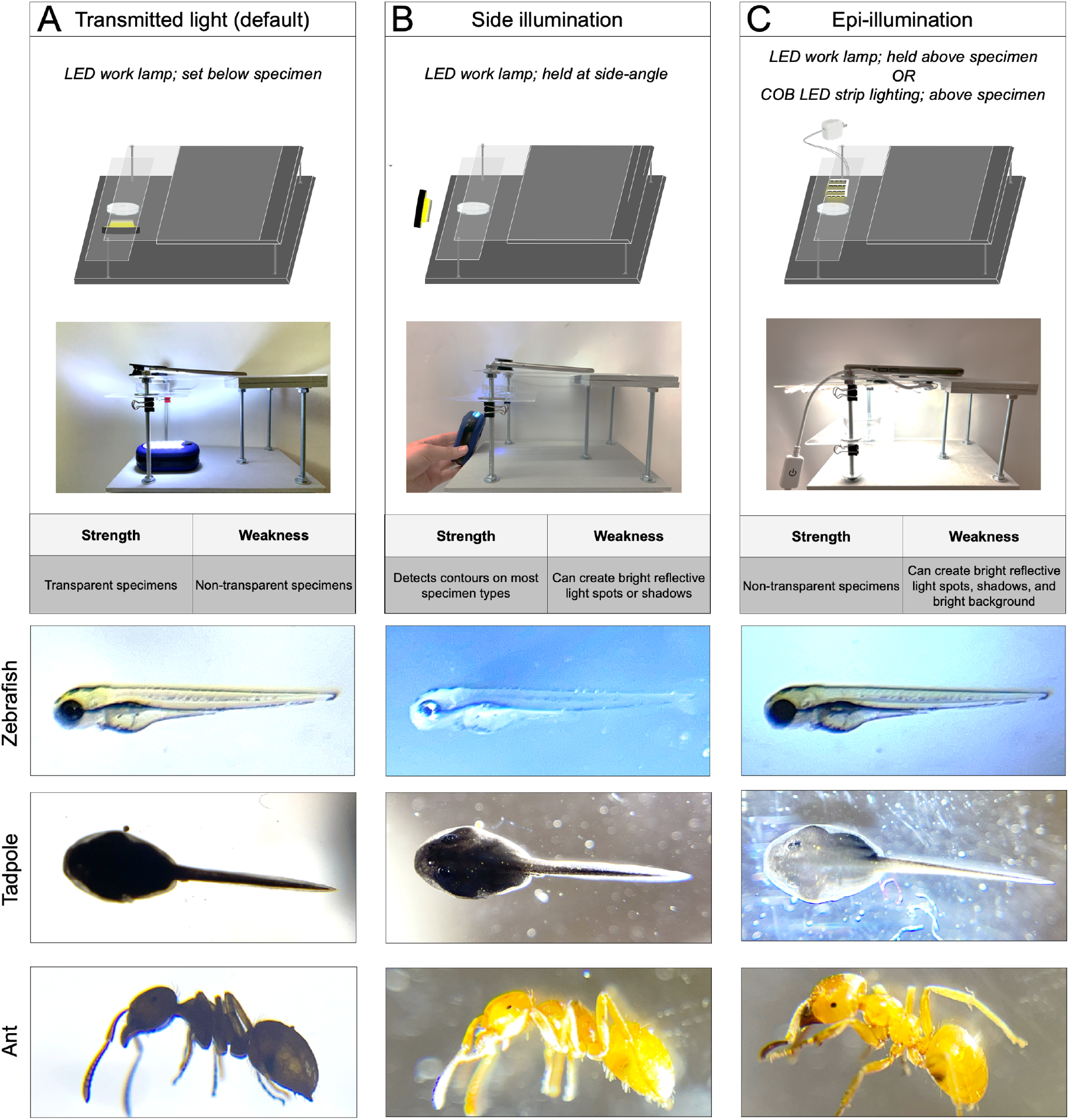
Optimized lighting for non-fluorescent viewing. Upper panels show the configuration, strengths, and weaknesses for (**A**) transmitted light, (**B**) side illumination, and (**C**) epi-illumination. Lower representative images show the same specimen viewed using each of the three lighting methods. Images shown were acquired using an Apple iPhone 12 Pro, 1x (middle magnification) lens, 6x digital zoom, with the additional clip-on macro lens.

### Use of glowscope devices for science education and outreach

Glowscopes may be suitable for some research applications, but our primary motivator for its development was to meet the needs of science educators and outreach programs. In the United States, recommendations for life science education at the K-12 and undergraduate levels involve increasing hands-on active learning ^14^. These recommendations deemphasize lecture-based information transfer as a sole means for student learning, and instead suggest that students learn science by iteratively performing the scientific method themselves. An inherent challenge of this pedagogical transformation is the increased pressure on teachers to develop new hands-on learning activities that are engaging and support the discipline-based learning goals. The glowscope may be one of many tools that is budget friendly, interests and excites students, and can support hands-on learning in classrooms. To aid its option, videos documenting its use and assembly instruction are available on a Glowscopes channel hosted on YouTube at https://www.youtube.com/channel/UCoRglCdvrtqwkBcvP10ibDg.

To assess potential opportunities for its use, we first examined the K-12 Next Generation Science Standards (NGSS). We identified NGSS disciplinary core ideas compatible with glowscope use at grade levels 1, 3, MS (middle school, grades 6-8), and HS (high school, grades 9-12), and developed a table of proposed student learning activities at each of these levels (Supplementary Tables S1-S3). Within this table, proposed student learning activities rely on glowscopes in different ways. For example, some activities use fluorescence while others use non-fluorescent viewing. Initial use of glowscopes without fluorescence may better prepare users for later fluorescence use, which is more technically challenging. Many activities propose use of zebrafish embryos, but other organisms can be used whether locally collected or purchased from biological suppliers. If use of zebrafish embryos is desired, educators should be aware of www.zfin.org as a means to search for nearby research labs already using zebrafish in their area, which may provide an opportunity for local access to obtain embryos or house adults.

We next performed a series of pilot experiments to provide representative examples of NGSS-compatible student learning activities for STEM education or outreach. In grade 1, NGSS recommends educators involve students in making observations as well as grade-appropriate proficiency in analyzing and interpreting data. We first focused on the specific core ideas addressing how offspring from the same parents can vary in appearance, and the types of behaviors that help offspring survive (Fig. 9). In these examples, students use non-fluorescent glowscopes to view animals on a smartphone or tablet display and develop tables to record their observations. To address how offspring behaviors help them to survive, we used a pencil to gently touch the embryos, observe their response, and construct tables for observations and the body parts involved in these observed responses. Although shown for tadpoles and monarch caterpillars (Fig. 9B), this activity would be compatible with zebrafish or many locally collected insects or worms.

**Figure 9.**
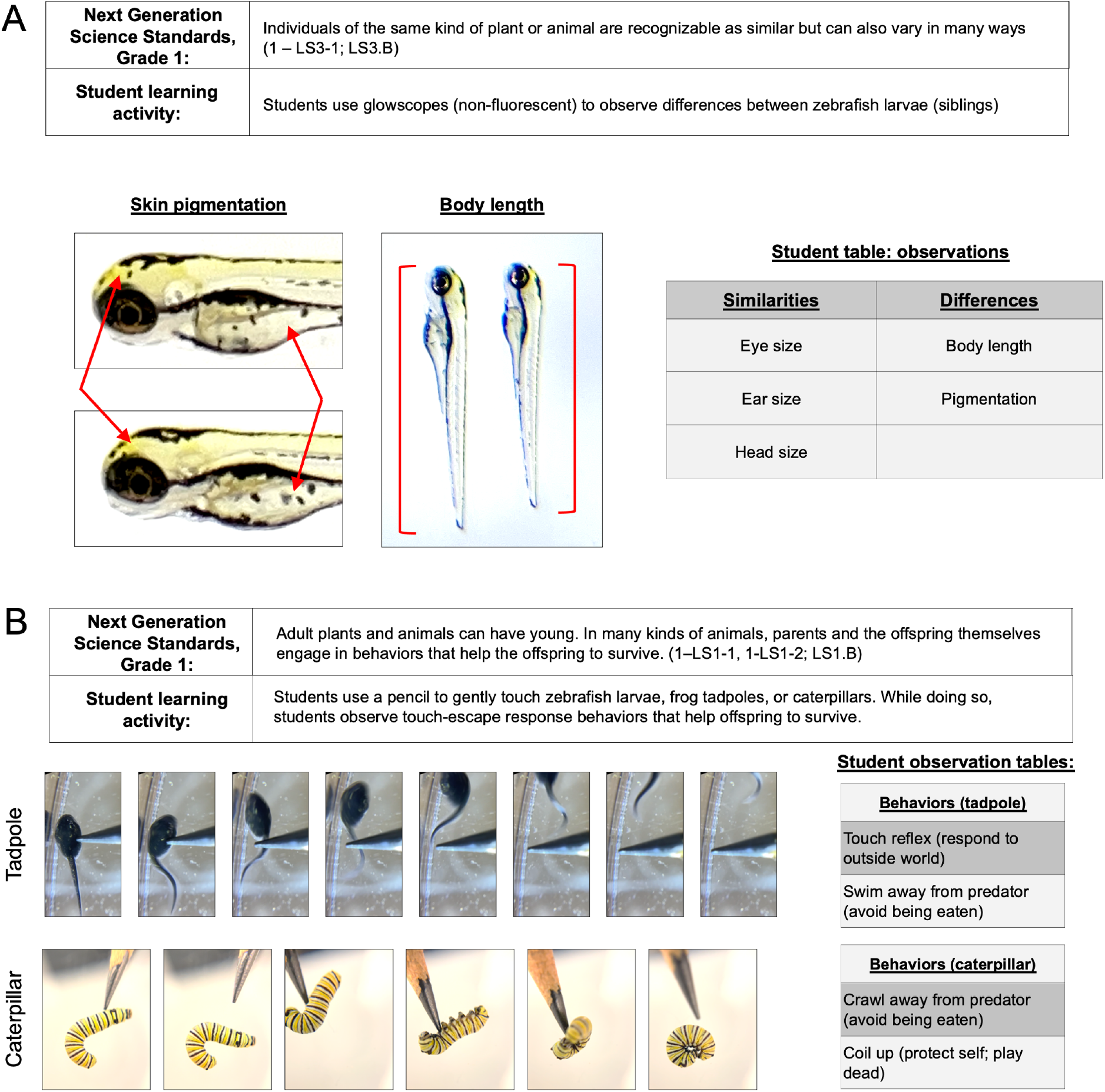
Grade 1 NGSS student learning activities using non-fluorescent glowscope. (**A**) Viewing of sibling zebrafish embryos to address how individuals of the same kind of animal are similar but can vary in many ways. Arrows on left point to skin pigment cells, which may represent melanocyotes, xanthophores, or iridophores. Note that their position and patterns on the body are variable from animal to animal. (**B**) Use of a pencil touch demonstrates offspring behaviors that aid their survival. Images from left to right are a time-series of the escape response. Images in A-B were acquired using an Apple iPhone 12 Pro.

An additional question addressed in the Grade 1 core ideas is how animals capture information needed for growth and respond in a way that helps them to survive (Fig. 10A). Zebrafish first obtain the ability to visually observe food, track and strike at moving food (prey), and use their jaw to eat food during their first week of life ^15,16^. Paramecia are a commonly used food source for young zebrafish. These single-celled organisms ‘swim’ in water using cilia to propel their movements. Individuals can be up to 0.5 mm in length and difficult to see with the naked eye, but their movements in petri dishes containing zebrafish larvae were easily observed using a smartphone with clip-on lens (Fig. 10B). Addition of paramecia to the petri dish containing zebrafish increased the frequency of swim events (Fig. 10C-E). We propose students compare non-fed with fed petri dishes (containing paramecia), and observe the types of behaviors and body parts used to respond to the presence of paramecia (Fig. 10F). These responses are essential for growth and survival and involve multiple organismal systems.

**Figure 10.**
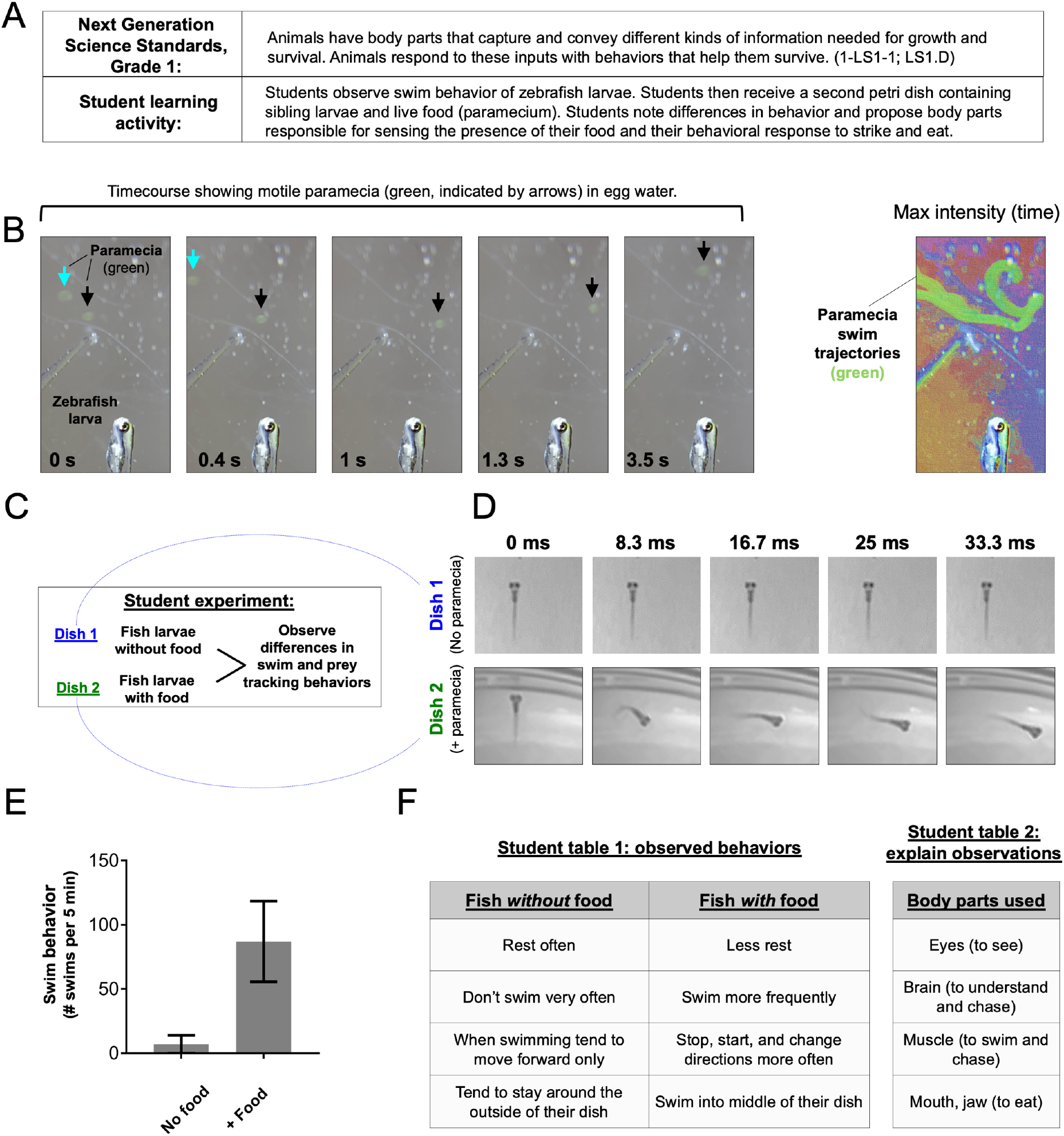
Grade 1 student learning activities addressing NGSS core ideas of the animal body parts needed for survival. (**A**) Description of NGSS standard and proposed student activity. (**B**) Timecourse (left, gray image series) shows two motile paramecia (blue and black arrows) in the same field of view as a paralyzed zebrafish larva (shown for scale comparison). Color image at right is a maximum intensity time projection, which plots the brightest pixel from each frame in a timelapse video and thereby marks the trajectory of each paramecia (green). Color was added to this image using the texturizer feature in PowerPoint. (**C**) Outline for student learning activity comparing swim and prey tracking behavior between a dish containing zebrafish larvae without food (paramecium) versus siblings in the presence of paramecia. (**D**) Timecourse image series shows zebrafish larva in the absence or presence of paramecia in the petri dish. Note the relative lack of movement in controls (upper, dish 1) in comparison to the paramecia treated dish (lower, dish 2). The timecourse shown in D represents 33 milliseconds. (**E**) Graph shows the difference in the frequency of fish larva (5 dpf) swim behavior between groups proposed in (D). N = 4 larvae per condition; bars show the mean ± standard error. (**F**) Example of student observations comparing groups from (D), and proposing the body parts involved in the prey tracking behaviors. Images were acquired using an Apple iPhone 12 Pro with (B) and without (D) the clip-on macro lens.

In high school life sciences (grades 9-12), NGSS core ideas focus on genes, inheritance mechanisms, and the effect of both genes and environment on the display of traits. To address the influence of environment on the display of traits, we developed a way to use glowscopes to monitor changes induced by temperature and water acidity. In these learning activities, students can investigate how individual zebrafish show changes to embryonic heart rate in response to these environmental stimuli (Fig. 11A-C). We demonstrate simple activities comparing heart rate before and after being placed onto a 35ºC reptile pad for 10 minutes (Fig. 11D). Fish may also experience changes to their environment in the form of water acidity. To model this, we added household vinegar to the water and observed decreased heart rate after 25-minute immersion in acidic water (Fig. 11D).

**Figure 11.**
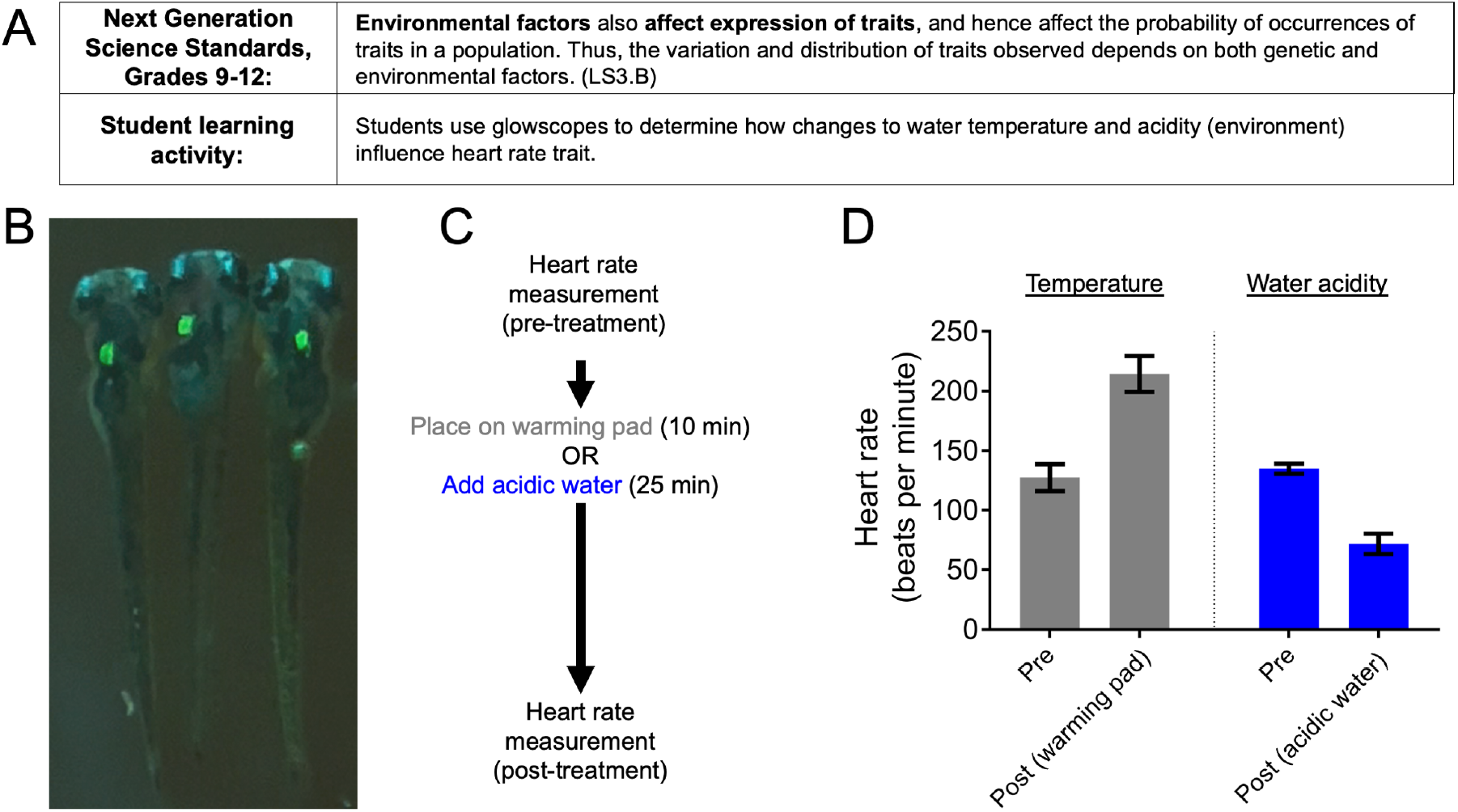
Grades 9-12 student learning activities addressing NGSS core ideas of how environment affects expression of traits. (**A**) Description of NGSS standard and proposed student activity. (**B**) Representative glowscope fluorescence image of 4 dpf *Tg(myl7:EGFP)* zebrafish larvae used for treatments and heart rate determination. (**C**) Experimental design for the influence of environment on heart rate trait. (**D**) Graph shows the effect of treatments on heart rate. Heart rate measurements on the same fish both prior to and after treatment, n = 4 larvae (temperature), n = 2 larvae (water acidity); bars show the mean ± standard deviation. Image shown in B was acquired using an Apple iPhone 12 Pro.

In addition to observing traits and their dependency on genetics and the environment, NGSS core ideas also emphasize mechanisms of genetic inheritance at the high school level (Fig. 12A). Because transgenes are dominant traits and are inherited in Mendelian patterns, the use of zebrafish ‘glow traits’ can be a fun and useful way for students to learn Punnett squares. In this learning activity, students are provided with several petri dishes, each containing embryos from different parental crosses (Fig. 12B). First, students observe each dish and the proportion of offspring displaying the ‘glow trait’. Students then develop Punnett squares for all possible parental genotypes, highlight which offspring genotypes will show dominant phenotypes, and then match their observed data to one or more Punnett squares. Students can then use their models to evaluate and discuss the possible parental genotypes for each dish of offspring.

**Figure 12.**
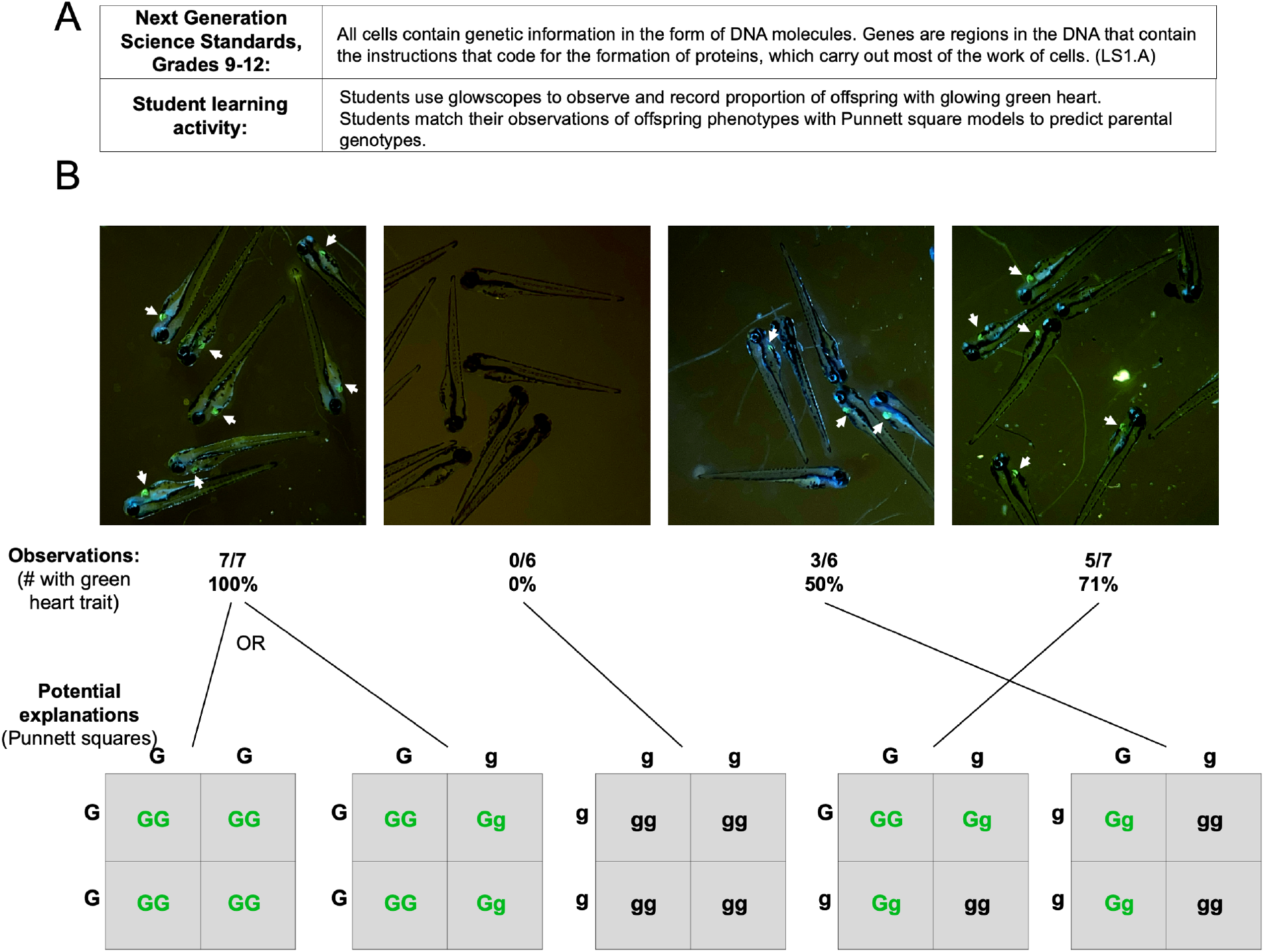
Grades 9-12 NGSS student learning activities focused on genetics and inheritance. (**A**) Description of NGSS standard and proposed student activity. (**B**) Representative glowscope fluorescence images of 4 dpf *Tg(myl7:EGFP)* zebrafish larvae used to determine the proportion inheriting the green heart trait (indicated by white arrows). Tabulated data are matched to Punnett squares consistent with the observed offspring phenotypic ratios. Images were acquired using an Apple iPhone 12 Pro.

## Discussion

In this study we developed and characterized components to convert a smartphone or tablet into a low-grade fluorescence microscope for a cost of $50 (US dollars) or less per unit. These build-it-yourself devices, which we refer to as glowscopes, are capable of imaging green and red fluorophores at up to 10 μm resolution and take advantage of the high frame rate capabilities of smartphone cameras for video recordings. The major advance of this study is the development of green and red fluorescence viewing using low-cost recreational LED flashlights and theater stage-lighting filters, which should be cross-compatible with microscope designs other than the smartphone and clip-on lens we used in our study. Due to the low cost, the specific design we characterize and report in this study should make it possible to equip entire classrooms with fluorescence imaging capabilities and student learning activities. In addition, we envision glowscopes could be useful for some research applications that do not require high magnification and have bright fluorescent specimens. For example, this may enable labs to concurrently acquire video data from many glowscopes at the same time, which may not be possible for labs that don’t have access to several fluorescence microscopes.

If used in combination with other low-cost or DIY smartphone microscopes such as Foldscopes, labs or students in classrooms can perform a range of microscopic images with tablets and smartphones. Specifically, Foldscopes can achieve at least five-fold greater magnification (exceeding 2 μm resolution) than the glowscope configuration we tested, making Foldscopes powerful devices for non-fluorescent smartphone imaging ^2^. This added resolution comes with tradeoffs that can be complemented by glowscopes. In addition to differences in fluorescence capability, the Foldscope’s added magnification has the tradeoff of reduced working distances, which can make it challenging or impossible to work with aquatic organisms or specimens in petri dishes. In comparison, the glowscope clip-on macro lens has longer and more flexible focal distances that make it compatible with large specimens and use of petri dishes, and can be used for non-fluorescent viewing with or without the clip-on macro lens. If used without the clip-on lens, the focal distance between the smartphone and specimen is on the order of 5-10 cm. In this way, the glowscope frame, with smartphone camera, can serve as a basic dissecting microscope.

Our testing in this study was performed using zebrafish embryos and larvae, but the smartphone microscope is not only compatible with zebrafish. However, if glowscope use is desired in K-12 or undergraduate education settings, zebrafish aquaria and research groups have become common to most colleges and universities and can be located using the ZFIN website to search for community members (people) by location (address; city name). This makes acquiring zebrafish embryos possible in most major cities. Many STEM outreach programs already exist that provide zebrafish to local educators, and in some instances, train educators ^17 18,19^. Additionally, with minimal instruction and equipment, educators could purchase adult GloFish from local retailers or online, perform fish breedings in small tanks, and acquire their own embryos as frequently as desired. Many transgenic reporter lines exist that express red and green fluorophores in diverse cell and tissue types and can be obtained through resource centers or by contacting laboratories directly. Based on our observations, any transgenic line expressing EGFP or red fluorophores would be compatible with glowscopes. Selecting those with brighter fluorescent reporter expression, more widespread expression (more cells and tissues), and with fluorescent proteins most closely matching the flashlight LED wavelength would be advantageous. For example, the ∼ 450 nm and 530 nm recreational LED flashlights we tested worked best with EGFP and DsRed, respectively.

Science education recommendations and evidence-based pedagogy has shifted away from the traditional lecture method and learner memorization of information. Instead, teaching and learning studies indicate that more active and inquiry-based learning engage students and support better learning outcomes ^20^. Providing students with opportunities to make their own observations, form models and predictions, acquire and interpret data, and use this to revise their models and understanding are common takeaways from life science education studies as well as NGSS and Vision and Change in Undergraduate Biology Education recommendations ^14^1. Use of devices such as glowscopes and Foldscopes can support these goals of allowing students to learn about science by doing science. In this way, students will build their own understanding of concepts rather than try to absorb information delivered in lecture format.

Our study primarily relied upon zebrafish for science education learning activities. Glowing transgenic zebrafish embryos or larvae offer students the chance to explore fundamental principles such as genes and proteins, inheritance, physiology, animal behavior, life cycles, and organismal development. In addition, zebrafish can also be a useful tool to address interactions between organisms and the environment due to the ease of adding factors to the embryo water in petri dish. For example, zebrafish heart structure and function, which can be easily viewed in the glowscope, is influenced by herbicides, pesticides, fungicides, alcohol, and tobacco ^21-25^. These and other recommendations listed in Supplementary Tables S1-S3 may provide a baseline for how educators can use glowscopes to excite learners of all ages, to match learning activities with NGSS disciplinary core ideas, and to address NGSS science and engineering practices.

## Availability of data and materials

All datasets are available upon request from the corresponding author.

## Funding

This work was supported by NSF CAREER award IOS-1845603 (J.H.H.) and Winona State University Foundation Special Projects award 251.0342 to Heather Nelson.

## Acknowledgements

We thank Erika Vail (Winona State University) for technical assistance, Carl Ferkinhoff (Winona State University) for advice and instrument sharing, and Amblynn Reisetter (Winona Public Schools) for helpful feedback.

## Notes

https://www.youtube.com/channel/UCoRglCdvrtqwkBcvP10ibDg

### Competing Interest Statement

The authors have declared no competing interest.

