## Supplementary Tables S1-S3 for "A low-cost smartphone fluorescence microscope for research, life science education, and STEM outreach"

### Supplementary Table S1

#### Next Generation Science Standards (NGSS) Primary; Grades K-5

| Grade Level | NGSS Disciplinary Core Ideas | | Potential student activities | Recommended viewing devices | NGSS Science and Engineering Practices | |
|---|---|---|---|---|---|---|
| K -- LS1-1 | Use observations to describe patterns of what plants and animals (including humans) need to survive. | Animals need food in order to live and grow (K-LS1-1). | Rear zebrafish (or tadpoles) to 1 month under different feeding conditions. **Observe** growth by comparing body length and size between groups. | None needed | **Scientific Knowledge is Based on Empirical Evidence** | Scientists look for patterns and order when **making observations** about the world. **Use observations to describe patterns** in the natural world in order to answer scientific questions. (K-LS1-1) |
| 1 – LS1-1 | Use materials to **design a solution** to a human problem by mimicking how plants and/or animals use their external parts to help them survive, grow, and meet their needs. | All organisms have external parts. Different animals use their body parts in different ways to see, hear, grasp objects, protect themselves, move from place to place, and seek, find, and take in food, water and air (LS1.A). | **Observe** zebrafish 3-7 day-old embryos or tadpoles and compare to insects. | Optional: Glowscope (without fluorescence) | **Constructing explanations and designing solutions** in K–2 builds on prior experiences and progresses to the use of evidence and ideas in constructing evidence-based accounts of natural phenomena and designing solutions. | **Make observations** (firsthand or from media) to **construct an evidence-based account** for natural phenomena. |
| | | Adult plants and animals can have young. In many kinds of animals, parents and the offspring themselves engage in behaviors that help the offspring to survive (LS1.B). Animals have body parts that capture and convey different kinds of information needed for growth and survival. Animals respond to these inputs with behaviors that help them survive (LS1.D). | **Observe** and touch zebrafish 3-7 day-old embryos or tadpoles with a hair-tool. **Construct an explanation** for how the observed behavior helps them to survive, and the body parts that capture and convey this information. | None needed | | |
| 1 – LS3-1 | **Make observations** to construct an evidence-based account that young plants and animals are like, but not exactly like, their parents. | Young animals are very much, but not exactly like, their parents. Individuals of the same kind of plant or animal are recognizable as similar but can also vary in many ways (LS3.A/B). | Use 3-7 day-old zebrafish embryos to **observe** any similar morphological features as well as potential variation in pigmentation traits. If available, include use of albino or casper mutant zebrafish strains. | Glowscope or Foldscope for high-magnification viewing of pigmentation | | |
| 3-LS3-1 | Analyze and interpret data to provide evidence that plants and animals have traits inherited from parents and that variation of these traits exists in a group of similar organisms. | Many characteristics of organisms are inherited from their parents (LS3.A). Different organisms vary in how they look and function because they have different inherited information (LS3.B). | Use 3-7 day-old zebrafish embryos to **observe** any similar morphological features as well as potential variation in pigmentation traits. If available, include use of albino or casper mutant zebrafish strains. | Glowscope or Foldscope for high-magnification viewing of pigmentation | **Analyzing data** in 3–5 builds on K–2 experiences and progresses to introducing quantitative approaches to collecting data and conducting multiple trials of qualitative observations. **Constructing explanations and designing solutions** in 3–5 builds on K–2 experiences and progresses to the use of evidence in constructing explanations that specify variables that describe and predict phenomena and in designing multiple solutions to design problems. | **Analyze and interpret data** to make sense of phenomena using logical reasoning. Use evidence (e.g., observations, patterns) to support an explanation. |
| 3-LS3-2 | Use evidence to support the explanation that traits can be influenced by the environment. | Other characteristics result from individuals' interactions with the environment, which can range from diet to learning. Many characteristics involve both inheritance and environment LS3.A). The environment also affects the traits that an organism develops (LS3.B). | **Observe** the body plan (morphology) and/or heart rate of 2-4 day-old zebrafish embryos. Consider altering the environment (egg water) by adding herbicide, pesticide, or fungicides (see Discussion). **Acquire evidence** to support an explanation for how the environment can influence traits. | Viewing of fluorescent (glowscope) or non-fluorescent heart (glowscope or Foldscope) | | |
| 3-LS4-3/4 | Construct an argument with evidence that in a particular habitat some organisms can survive well, some survive less well, and some cannot survive at all. Make a claim about the merit of a solution to a problem caused when the environment changes and the types of plants and animals that live there may change.* | When the environment changes in ways that affect a place's physical characteristics, temperature, or availability of resources, some organisms survive and reproduce, others move to new locations, yet others move into the transformed environment, and some die (LS2.C). For any particular environment, some kinds of organisms survive well, some survive less well, and some cannot survive at all. (LS4.C). Populations live in a variety of habitats, and change in those habitats affects the organisms living there (LS4.D). | **Observe** the physical characteristics (growth, swim behavior, heart rate) and survival of 1-10 day-old zebrafish embryos. Consider altering the environment (embryo water) by altering the pH or temperature. **Acquire evidence** and **analyze data** to support an **explanation** for how the environment can influence traits and survival. | None needed. (Optional): view heart rate using glowscope | | |

#### Supplementary Table S2

**Next Generation Science Standards (NGSS)** **Intermediate; Middle School Level**

| Grade Level | NGSS Disciplinary Core Ideas | | Potential student learning activities | Recommended viewing devices | NGSS Science and Engineering Practices | |
|---|---|---|---|---|---|---|
| MS-LS1-1 | Conduct an investigation to provide evidence that living things are made of cells; either one cell or many different numbers and types of cells. | All living things are made up of cells, which is the smallest unit that can be said to be alive. An organism may consist of one single cell (unicellular) or many different numbers and types of cells (multicellular) (LS1.A). | **Construct a model** for how specialized cells with pigment granules, if spatially localized and distributed throughout the animal, could be responsible for organismal trait like stripes and coloration. Observe pigment cells on the 3-7 day-old zebrafish embryo to **revise model.** | Recommended: Foldscope or glowscope (non-fluorescent) | **Developing and using models**. Modeling in 6–8 builds on K–5 experiences and progresses to developing, using, and revising models to describe, test, and predict more abstract phenomena and design systems. **Planning and Carrying out Investigations** in 6-8 builds on K- 5 experiences and progresses to include investigations that use multiple variables and provide evidence to support explanations or solutions. **Constructing Explanations and Designing Solutions** in 6–8 builds on K–5 experiences and progresses to include constructing explanations and designing solutions supported by multiple sources of evidence consistent with scientific knowledge, principles, and theories. **Engaging in argument from evidence.** Engaging in argument from evidence in 6–8 builds on K–5 experiences and progresses to constructing a convincing argument that supports or refutes claims for either explanations or solutions about the natural and designed world(s). | **Develop and use a model** to describe phenomena. AND Develop a model to describe unobservable mechanisms. **Conduct an investigation** to produce data to serve as the basis for evidence that meet the goals of an investigation. **Construct a scientific explanation based on valid and reliable evidence** obtained from sources (including the students' own experiments) and the assumption that theories and laws that describe the natural world operate today as they did in the past and will continue to do so in the future. **Use an oral and written argument supported by evidence** to support or refute an explanation or a model for a phenomenon. (MS-LS1-3)<br>▪ Use an oral and written argument supported by empirical evidence and scientific reasoning to support or refute an explanation or a model for a phenomenon or a **solution to a problem.** (MS-LS1-4) |
| MS-LS1-3 | Use argument supported by evidence for how the body is a system of interacting subsystems composed of groups of cells. | In multicellular organisms, the body is a system of multiple interacting subsystems. These subsystems are groups of cells that work together to form tissues and organs that are specialized for particular body functions (LS1.A). | **Construct a model** for the specialized cells that form heart chambers and valves that are specialized for the functions they provide, and how these groups of cells behave differently than most somatic cell types (contractility). Observe the beating heart using the glowscope. **Construct an explanation** for how these structures and cell behaviors confer the function of the system (blood flow that is unidirectional). **Plan and carry out an investigation** using high dose anesthetic treatment (tricaine) to block observed heart contractility and draw comparisons of red blood cell movement through vasculature between groups. **Use evidence** to support or reject earlier models. | Recommended: Glowscope viewing of fluorescent heart. Foldscope viewing of blood cell movement through vasculature. | | |
| MS-LS1-5 | Construct a scientific explanation based on evidence for how environmental and genetic factors influence the growth of organisms. | Genetic factors as well as local conditions affect the growth of the adult plant or animal (LS1.B). | **Plan and carry out an investigation** to determine how the density of zebrafish (per dish) influences their growth. **Interpret evidence** and **construct an argument** to refute or support the claim that environment influences animal growth. | Recommended: Glowscope (non-fluorescent) and length measurements. | | |
| MS-LS1-8 | Gather and synthesize information that sensory receptors respond to stimuli by sending messages to the brain for immediate behavior or storage as memories. | Each sense receptor responds to different inputs (electromagnetic, mechanical, chemical), transmitting them as signals that travel along nerve cells to the brain. The signals are then processed in the brain, resulting in immediate behaviors or memories (LS1.D). | **Construct a model** for how sensory information (touch) is relayed to the brain. **Plan and carry out an investigation** to determine if 5-10 day-old zebrafish larvae learn to reduce their sensitivity and response to repeated touch, increase their swim behavior in response to the presence of food (paramecium), or decrease their sensitivity after being continuously provided food for a day or more.. **Interpret evidence** and **construct an argument** to refute or support the claim that the brain interprets touch sensation. | None needed | | |
| MS-LS3-1 | Develop and use a model to describe why structural changes to genes (mutations) located on chromosomes may affect proteins and may result in harmful, beneficial, or neutral effects to the structure and function of the organism. | Genes are located in the chromosomes of cells, with each chromosome pair containing two variants of each of many distinct genes. Each distinct gene chiefly controls the production of specific proteins, which in turn affects the traits of the individual. Changes (mutations) to genes can result in changes to proteins, which can affect the structures and functions of the organism and thereby change traits (LS3.A). In addition to variations that arise from sexual reproduction, genetic information can be altered because of mutations. Though rare, mutations may result in changes to the structure and function of proteins. Some changes are beneficial, others harmful, and some neutral to the organism (LS3.B). | **Observe** transgenic zebrafish and non-transgenic siblings. **Construct a model** for why some but not all display the phenotypic trait. After learning about the green/red fluorescent protein gene and protein, **Plan and carry out an investigation** to determine if may be genetically inherited. **Use evidence** to support or reject earlier models and predictions. **Construct an argument** that the trait is encoded by a heritable gene. | Glowscope (fluorescent) | | |
| MS-LS3-1, 2 | Develop and use a model to describe why asexual reproduction results in offspring with identical genetic information and sexual reproduction results in offspring with genetic variation. | Organisms reproduce, either sexually or asexually, and transfer their genetic information to their offspring (LS1.B). Variations of inherited traits between parent and offspring arise from genetic differences that result from the subset of chromosomes (and therefore genes) inherited (LS3.A). In sexually reproducing organisms, each parent contributes half of the genes acquired (at random) by the offspring. Individuals have two of each chromosome and hence two alleles of each gene, one acquired from each parent. These versions may be identical or may differ from each other (LS3.B). | | | | |

#### Supplementary Table S3

**Next Generation Science Standards (NGSS)** | **Secondary; High School Level**

| **Grade Level** | **NGSS Disciplinary Core Ideas** | | **Potential student learning activities** | **Recommended viewing devices** | **NGSS Science and Engineering Practices** | |
|---|---|---|---|---|---|---|
| HS-LS1-1 | Construct an explanation based on evidence for how the structure of DNA determines the structure of proteins, which carry out the essential functions of life through systems of specialized cells. | All cells contain genetic information in the form of DNA molecules. Genes are regions in the DNA that contain the instructions that code for the formation of proteins, which carry out most of the work of cells (LS1.A). | **Develop a model** of genes, chromosomes, and DNA and their relatedness to proteins within cells. **Observe** zebrafish offspring from adults (various parental crosses) that produce varying ratios of offspring possessing the trait. Determine the proportion carrying the trait and match this to possible Punnett squares. More advanced: **Observe** zebrafish offspring from adults that carried the (trans)genes for green and red fluorescent traits. **Plan and carry out an investigation,** using **mathematical analysis**, for the ratio of offspring displaying one or both traits and whether they are linked. **Use evidence to revise models**, if needed. | Glowscope (green and red fluorescence) | **Developing and Using Models** Modeling in 9–12 builds on K–8 experiences and progresses to using, synthesizing, and developing models to predict and show relationships among variables between systems and their components in the natural and designed worlds. **Planning and Carrying Out Investigations in** 9-12 builds on K-8 experiences and progresses to include investigations that provide evidence for and test conceptual, mathematical, physical, and empirical models. **Constructing explanations and designing solutions** in 9–12 builds on K–8 experiences and progresses to explanations and designs that are supported by multiple and independent student- generated sources of evidence consistent with scientific ideas, principles, and theories. | **Develop and use a model** based on evidence to illustrate the relationships between systems or between components of a system. (HS-LS1-2), Use a model based on evidence to illustrate the relationships between systems or between components of a system. **Plan and conduct an investigation** individually and collaboratively to produce data to serve as the basis for evidence, and in the design: decide on types, how much, and accuracy of data needed to produce reliable measurements and consider limitations on the precision of the data (e.g., number of trials, cost, risk, time), and refine the design accordingly. **Construct an explanation** based on valid and reliable evidence obtained from a variety of sources (including students' own investigations, models, theories, simulations, peer review) and the assumption that theories and laws that describe the natural world operate today as they did in the past and will continue to do so in the future. (HS-LS1-1) **Construct and revise an explanation** based on valid and reliable evidence obtained from a variety of sources (including students' own investigations, models, theories, simulations, peer review) and the assumption that theories and laws that describe the natural world operate today as they did in the past and will continue to do so in the future. (HS-LS1-6) |
| HS-LS1-4 | Use a model to illustrate the role of cellular division (mitosis) and differentiation in producing and maintaining complex organisms. | In multicellular organisms individual cells grow and then divide via a process called mitosis, thereby allowing the organism to grow. The organism begins as a single cell (fertilized egg) that divides successively to produce many cells, with each parent cell passing identical genetic material (two variants of each chromosome pair) to both daughter cells. Cellular division and differentiation produce and maintain a complex organism, composed of systems of tissues and organs that work together to meet the needs of the whole organism (LS1.B). | **Observe** early embryonic development 1-2-4-8 cell stage divisions of zebrafish eggs (or frog, sea urchin, or others available. **Develop a model** for the genetic material and whether/how it is passed from parental cell(s) to daughter cells within the organism. | Foldscope (or glowscope), non-fluorescent | | |
| HS-LS3-2/3; HS-LS2-7 | Make and defend a claim based on evidence that inheritable genetic variations may result from (1) new genetic combinations through meiosis, (2) viable errors occurring during replication, and/or (3) mutations caused by environmental factors AND Apply concepts of statistics and probability to explain the variation and distribution of expressed traits in a population. Design, evaluate, and refine a solution for reducing the impacts of human activities on the environment and biodiversity. | Environmental factors can also cause mutations in genes, and viable mutations are inherited. Environmental factors also affect expression of traits, and hence affect the probability of occurrences of traits in a population. Thus the variation and distribution of traits observed depends on both genetic and environmental factors (LS3.B). When evaluating solutions it is important to take into account a range of constraints including cost, safety, reliability and aesthetics and to consider social, cultural and environmental impacts (ETS1.B). | **Design and investigation** to test how environmental factors such as herbicides, pesticides, fungicides, drugs such as allergy pills with adverse side effects, or waste pharmaceuticals influence the display of traits (zebrafish heart rate). **Use evidence** to **design a solution** based on observed data and other constraints including cost, effects on agriculture, safety, social, cultural, and environmental impacts. | Glowscope | | |
