## Supplementary Methods for "A low-cost smartphone fluorescence microscope for research, life science education, and STEM outreach"

### Wood or composite board frame preparation

1. Plywood or composite board lower frame. Cut to 7.5” x 10.5”. Drill five holes with a ¼” drill bit based on the design coordinates below.
2. Plywood or composite board upper pieces (these support the plexiglass top). Cut two pieces to 7.5” x 4”.
    a. In the first upper piece, which will eventually be the middle piece, drill with a ¼” drill bit (design coordinates below, lower left image).
    b. In the second upper piece, which will eventually be the top piece, drill holes (design coordinates below, lower right image) using a 5/8” drill bit.

**the center of each drilled hole should be 1” from each side.*

**Design coordinates for frame preparation**

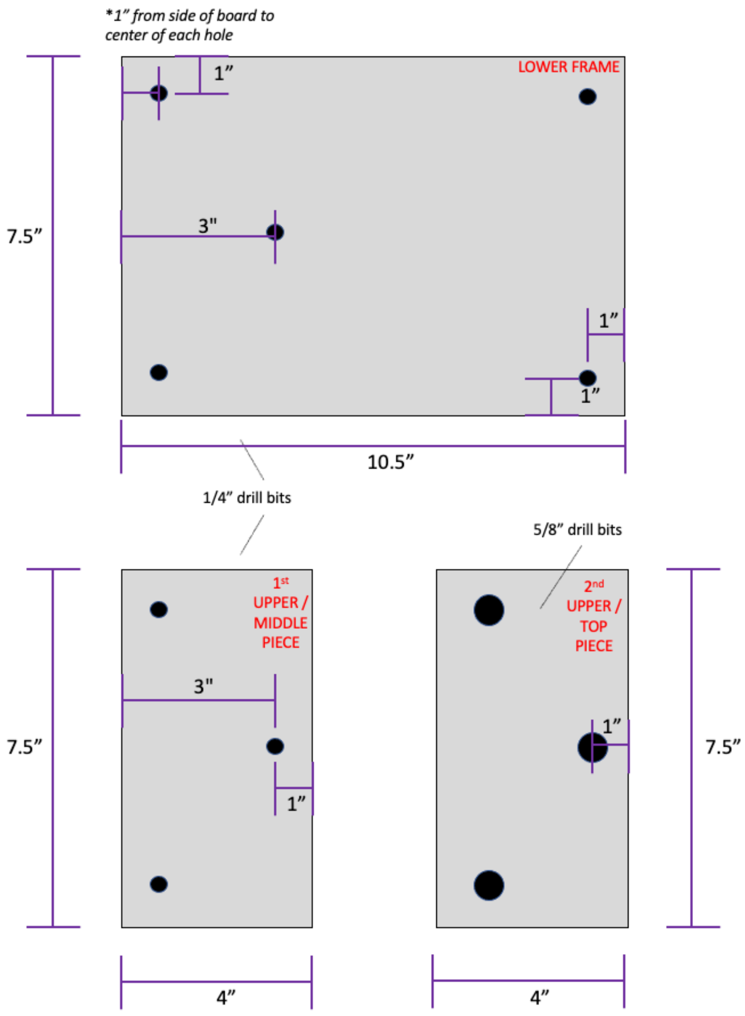

#### Plexiglass preparation

*Use of an oscillating tool with plastic or wood compatible blade is best. Alternatively, a glass or plexiglass scoring tool (sharp like a knife), when used along with clamps and straightedge, can be used to score and then snap the plexiglass.*

*After cutting or snapping, a bench grinder (grinding wheel) can be useful to smooth imperfections or round sharp corners.*

3. Upper large platform. Cut or snap to 7.5” x 9”. Drill the two corner holes using the design coordinates below using a ¼” bit. Then, drill the desired number and size of camera viewing holes between these two corner holes. See images.

   *Use of a step drill bit (1/4” – ¾” cone shaped bit”) can be useful to open up the viewing holes in a way that will look perfectly circular and smooth.*

4. Lower stage platform. Cut or snap to 7.5” x 4”. Drill two holes using the design coordinates below using a ¼” bit.

**Design coordinates for plexiglass preparation**

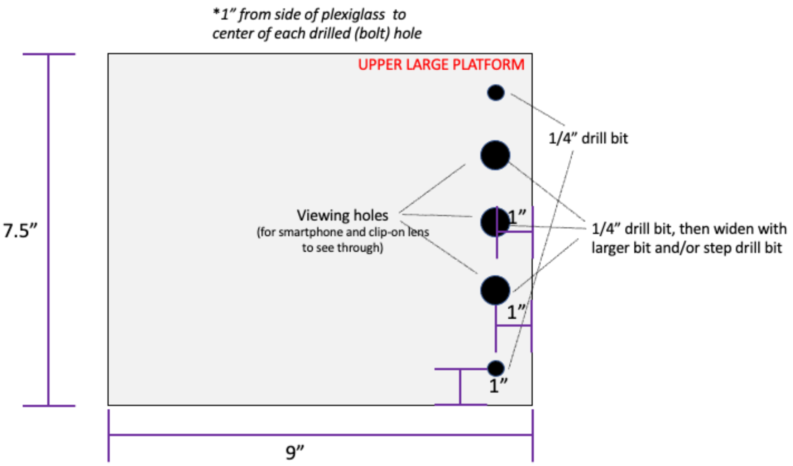


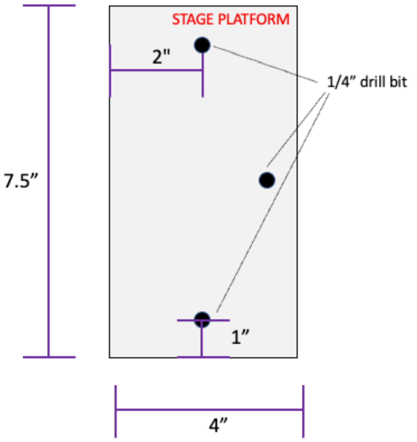

### Assembly

5. Assemble the frame parts based on the illustration below.

**Use of a ratcheting wrench made accelerates assembly*

#### Assembly illustration

##### A. Assemble frame base

1. Insert carriage bolts through holes in board
2. Per bolt: one standard ¼" washer and ¼" hex nut, tighten

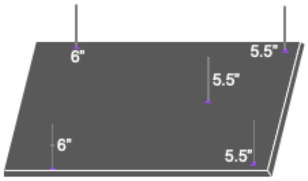


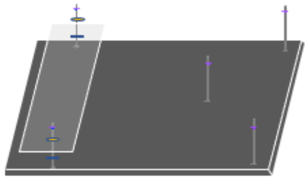

##### B. Attach stage, prep for top pieces

1. ¼" Fender washer (below stage)
2. Plexiglass stage piece
3. ¼" Fender washer (above stage)
4. Standard ¼" hex nut near top of each 5.5" bolt
5. ¼" jam nut (low profile nut) near top of each 6" bolt
6. Standard 1/4" washer above each nut

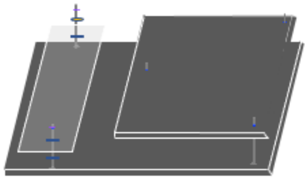

##### C. Attach upper middle piece

1. Add 'upper middle piece' (wood/composite)
2. Add a **¼" jam nut** above the piece onto each of the three 5.5" bolt. Tighten down.

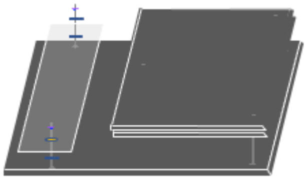

##### D. Add upper top piece

1. Add 'upper top piece' (wood/composite).

**This piece just rests atop and jam nuts should be even with or just lower than the top of this 'upper top piece'. The jam nut and carriage bolt end should not protrude above this 'upper top piece' or they will scratch and break the plexiglass in next steps.*

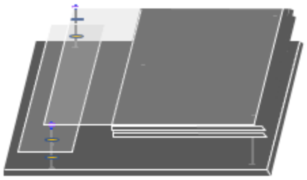

##### E. Add, adjust, and tighten down plexiglass upper large platform

1. Insert pre-drilled **plexiglass platform** (2 holes) over and onto the two 6" carriage bolts (above stage)
2. Add a standard ¼" washer on each side
3. Adjust the nuts below plexiglass platform, along with the wooden platform, to make the upper plexiglass platform as level as possible.
4. Add a standard ¼" hex nut onto the top of each 6" carriage bolt (above plexiglass and washer). Tighten gently. (over-tightening may crack the plexiglass).

#### Stage positioning and use

6. Slide the stage up and down to the desired position. The use of fender washers and clamps both above and below the stage holds it into position and reduces its wobble. This will reduce the frequency that the specimen / dish slides or falls off the stage. See the illustrations below.

   *We've found that when using the extra small binder clamps (15 mm size) and a hex jam nut (low profile) underneath the upper plexiglass platform, pushing the stage as high as possible works perfectly with the clip-on macro lenses we tested.*

**<u>Stage positioning illustration</u>**

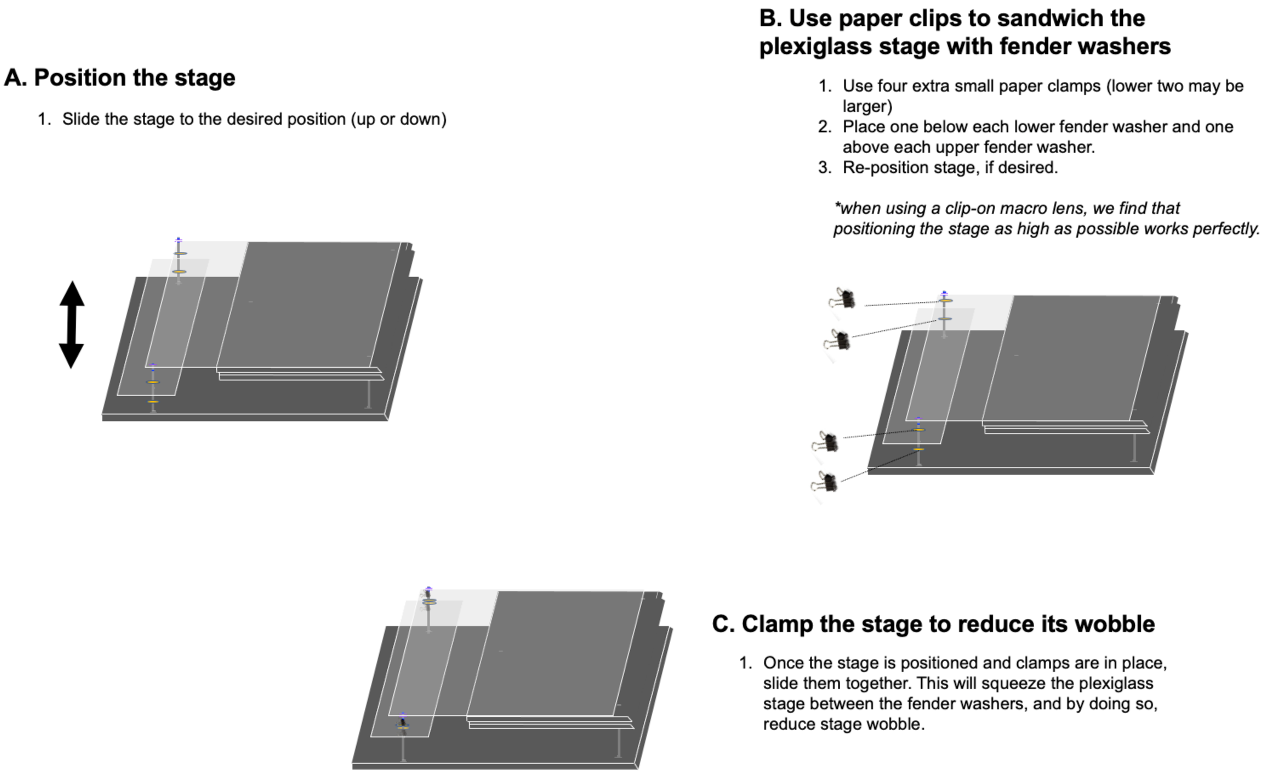
